# From Scalp to Source: Precise Phase Retrieval of Intracerebral Epileptic Sources Based on Surface EEG

**DOI:** 10.64898/2026.08.16.745081

**Authors:** Kristóf Furuglyás, Márton Huszár-Kis, Bálint Horváth, Andrea Pejin, Nóra Forgó, István Langó, Shobhit Singla, Márton Görög, Péter Vass, Zoltán Chadaide, Tamás Laszlovszky, Orrin Devinsky, Anto I. Bagić, Zoltán Somogyvári, Antal Berényi

## Abstract

Accurate phase tracking of deep-brain activity is critical for effective closed-loop and phase-locked neuromodulation therapies. However, direct access to deep neural phase through intracranial recordings remains clinically restrictive due to the invasiveness. Here we validate and clinically benchmark the Gábor-Nelson (GN) dipole estimation method for reconstructing deep-brain oscillatory phase from non-invasive scalp EEG. GN is a geometry-based, imaging-independent approach that offers computationally efficient dipole reconstruction and has rarely been applied to source-level phase estimation in human neuroscience. We compared GN with an established MRI-informed Inverse Solution (IS) method using a three-stage reconstruction pipeline consisting of dipole modeling, dimensionality reduction, and frequency-dependent phase-delay correction. Validation is performed using (i) cadaveric recordings, where known ground-truth seizure waveforms were replayed through implanted deep electrodes, and (ii) simultaneous scalp EEG and SEEG recordings in human patients, where pseudo-ground truth was approximated via the intracranial contacts. GN achieved phase accuracy and signal fidelity comparable to IS across both datasets despite requiring no anatomical imaging. In cadaver recordings, phase-corrected reconstruction correlations exceeded r > 0.91 and Δφ < 9° in mean phase error. In patient SEEG data, GN reached up to r ≈ 0.80 with phase offsets suitable for neuromodulatory timing. GN offers a viable, low-barrier, imaging-independent alternative to traditional inverse modeling for non-invasive seizure phase tracking. This framework opens pathways for scalable, phase-locked and closed-loop stimulation therapies in epilepsy and potentially other network-based brain disorders.

## 1 Introduction

Accurately tracking the instantaneous phase of neural oscillations within deep-brain sources is a major prerequisite for the success of next-generation phase-locked neuromodulation therapies. Because the clinical efficacy of closed-loop electrical stimulation relies on subsecond-level precision, stimulation pulses must be synchronized with specific, ongoing oscillatory states of the underlying neural networks. Phase-specific interventions have already demonstrated remarkable therapeutic potential, showing capability to terminate epileptic seizures [1,2], alleviate depression-like symptoms [3], and enhance fear extinction pathways [4]. Recently, Guo et al. [5] demonstrated that cycle-by-cycle phase-locked deep brain stimulation can dynamically modulate both pathological neural activity and emergent motor behaviors in Parkinson’s disease. Furthermore, translational computational models indicate that delayed feedback protocols can actively suppress pathological network synchrony in movement disorders [6]. Because these adaptive control strategies depend on the temporal fidelity of the underlying oscillations, achieving real-time phase tracking remains a paramount engineering challenge.

Directly capturing deep-brain oscillatory dynamics via intracranial recordings remains clinically constrained due to the inherent surgical risks and restricted accessibility of penetrating depth electrodes. This clinical bottleneck highlights an urgent need for non-invasive frameworks capable of accurately reconstructing deep-source phase trajectories from surface scalp EEG recordings. While several recent studies confirm that signals originating from hidden or mesial structures, such as the hippocampus, amygdala, insula, and orbitofrontal cortices, propagate detectable signatures to scalp EEG and MEG sensors [7–11], non-invasive isolation remains exceptionally difficult. Highly attenuated deep signals are severely masked by folded cortical anatomy, signal cancellation, and prominent, superimposed neocortical background activity [10]. Consequently, while deep epileptic activity is partially accessible from the scalp, extracting clean, temporally high-fidelity waveforms requires advanced, computationally heavy reconstruction methodologies to achieve clinical utility.

In a recent review, Santos et al. (2025) highlighted that while concurrent scalp-SEEG recordings offer complementary insights into epileptic networks, historical investigations focused almost exclusively on seizure detection, electrographic pattern correspondence, and spatial localization, rather than source-level phase reconstruction [12]. Similarly, parallel invasive and non-invasive recording configurations have primarily served as static ground-truth benchmarks for validating classical spatial source-localization accuracy. This includes Pigorini et al. (2024) linking macro-scale scalp signals to micro-scale intracranial electrophysiology [13], and Mikulan et al. (2020) utilizing high-density EEG to map intracerebral stimulation responses. While Parmigiani et al. (2022) utilized cortico-cortical evoked potentials (CCEPs) to demonstrate that scalp profiles can reliably describe deep electrical stimulation effects [14], and Subramanian et al. (2025) proved that scalp arrays can predict lower-frequency intracranial activity [7], these approaches rarely address the temporal distortions corrupting the signal phase.

Unlike direct stereo-EEG (SEEG) recordings, surface measurements suffer from spatial blurring and phase distortion induced by volume conduction, low-pass tissue filtering, and source–sensor misalignment [7]. Furthermore, non-ideal hardware transfer function, and capacitive filtering at the electrode–tissue interface introduce frequency-dependent phase-shifts and signal dispersion [15]. Given, that the scalp EEG phase carries critical information regarding the underlying multi-unit spiking activity (MUA) [16], correcting these timing distortions is essential. While traditional source-imaging prioritizes delineating the anatomical boundaries of the seizure onset zone, the primary objective of this study is fundamentally different: we ask whether the dominant phase of the deep epileptic generator can be reconstructed with the precise temporal resolution required for phase-locked neuromodulation, bypassing the computationally heavy full-localization problem entirely.

To resolve these propagation issues, conventional EEG source-localization relies heavily on distributed inverse solutions or sparse Bayesian approaches [17–19]. Although mathematically rigorous, these frameworks require subject specific anatomical imaging, precise electrode co-registration, realistic head models, and extensive parameter tuning, which collectively limit their scalability in routine clinical workflows. In equivalent-current-dipole (ECD) modeling, localization is formulated as an optimization problem where dipole positions, orientations, and moments are iteratively fitted to minimize the residual variance between predicted and scalp-recorded topographies [20]. Extending this to multi-dipole configurations severely expands the parameter space, introducing highly non-linear, high-dimensional optimization challenges [21,22]. Crucially, as noted by Bastola et al. (2024), low signal-to-noise rations (SNR) and deep seated generators often cause these non-linear optimization algorithms to stall in local minima or converge onto erroneous anatomical locations [23]. This risk underscores the clinical value of evaluating a simpler, imaging-independent solution, when high-resolution structural mapping is secondary to rapid, temporally precise phase tracking.

To address this gap, we introduce the Gábor–Nelson method, a geometry-based, model-light mathematical approach optimized for phase reconstruction [24,25], and benchmark its performance against a rigorous, MRI-informed Inverse Solution (IS) pipeline. While the benchmark IS method relies on subject-specific forward models of volume conduction, the GN method acts as an anatomy-independent estimator utilizing surface integral mathematics. To ensure standardized comparison, both algorithms are embedded into a uniform three-stage processing pipeline: (1) Dipole Modeling: Reconstructing time-varying three-dimensional dipole moment vectors from multi-channel surface EEG signals; (2) Dimensionality Reduction: Applying independent component analysis (ICA) to isolate the single, dominant oscillatory phase component from the 3D trajectory; and (3) Phase-Delay Correction: Implementing a frequency-dependent correction function to compensate for hardware and biophysical propagation delays.

We validate this comprehensive framework across two distinct reference datasets. First, we establish absolute technical precision using controlled human cadaver recordings, where known, pre-recorded seizure waveforms were physically replayed through implanted deep electrodes into human cadaver heads to mimic deep-brain signal sources, allowing direct benchmarking against known ground truth. Second, we assess in vivo clinical utility using simultaneous scalp–SEEG recordings from patients with focal epilepsy, approximating a pseudo-ground truth directly from intracranial contacts. This dual-validation paradigm effectively dissociates spatial localization errors from temporal phase reconstruction accuracy. Ultimately, by evaluating signal fidelity and circular phase alignment across both datasets, we demonstrate that deep-brain epileptic phase dynamics can be non-invasively recovered with the precision required to guide future closed-loop neuromodulation systems.

## 2 Materials and Methods

This section describes the complete methodological framework developed to reconstruct the phase of deep epileptic sources from surface EEG recordings.

### 2.1 Overview of the Phase Reconstruction Pipeline

This framework was applied uniformly across both cadaveric and SEEG datasets, with adaptations in preprocessing and ground truth estimation where necessary.

In the first stage, we estimate the time-varying dipole moment that best accounts for the observed scalp EEG. We implemented two complementary methods. A geometry-based, anatomy-independent method relying on surface potentials and electrode positioning (i.e., the ‘Gábor-Nelson’ (GN) method), and a model-based approach leveraging subject-specific MRI data and forward modeling of volume conduction (i.e., the ‘Inverse Solution’ (IS) method). Both methods yield a three-dimensional dipole vector at each time point, capturing the temporal evolution of source activity in Cartesian space.

Next, to extract a unidimensional signal suitable for phase analysis, Independent Component Analysis (ICA) is applied to the reconstructed 3D dipole time series. This step isolates the dominant oscillatory component and enables continuous phase assignment over time. The source separation capability of ICA was critical for isolating the true oscillatory component of interest [26]. This strategy ensured that the phase analysis was always performed on a signal that best represented the underlying epileptic activity, improving the fidelity of our phase-based analysis of early focal seizures. The approach is consistent with general best practices in signal processing for epilepsy i.e. to use ICA when dealing with multiple concurrent signals or noise [26–28].

Hardware-dependent phase shifts were addressed in the third step, specifically those occurring in cadaver experiments. These differences, arising from the interplay between the replaying and recording systems, introduced additional delays and distortions that required compensation. Recorded EEG signals are subject to frequency-dependent phase shifts introduced by the hardware’s non-ideal transfer function and biophysical filtering effects. To correct these distortions, we computed a frequency-domain phase-delay correction function using Fourier spectrum analysis. The comparison between the replayed known ground truth signal and the recorded one allowed us to estimate the phase distortion introduced by the hardware. No phase correction was applied to SEEG datasets, as both deep and surface recordings were acquired using the same amplifier system, thereby eliminating inter-modality hardware-related phase discrepancies.

Lastly, for SEEG datasets, where direct ground truth is unavailable, we established a pseudo-ground truth derived from deep recordings. This was typically approximated using spatial derivatives of intracranial recordings (e.g., 1D current source density) and deep dipole modeling, which served as reference signals for validating the reconstructed phase trajectories.

A schematic overview of the complete reconstruction and validation workflow, including dataset-specific ground truth integration, is shown in Figure 1.

**Figure 1.**
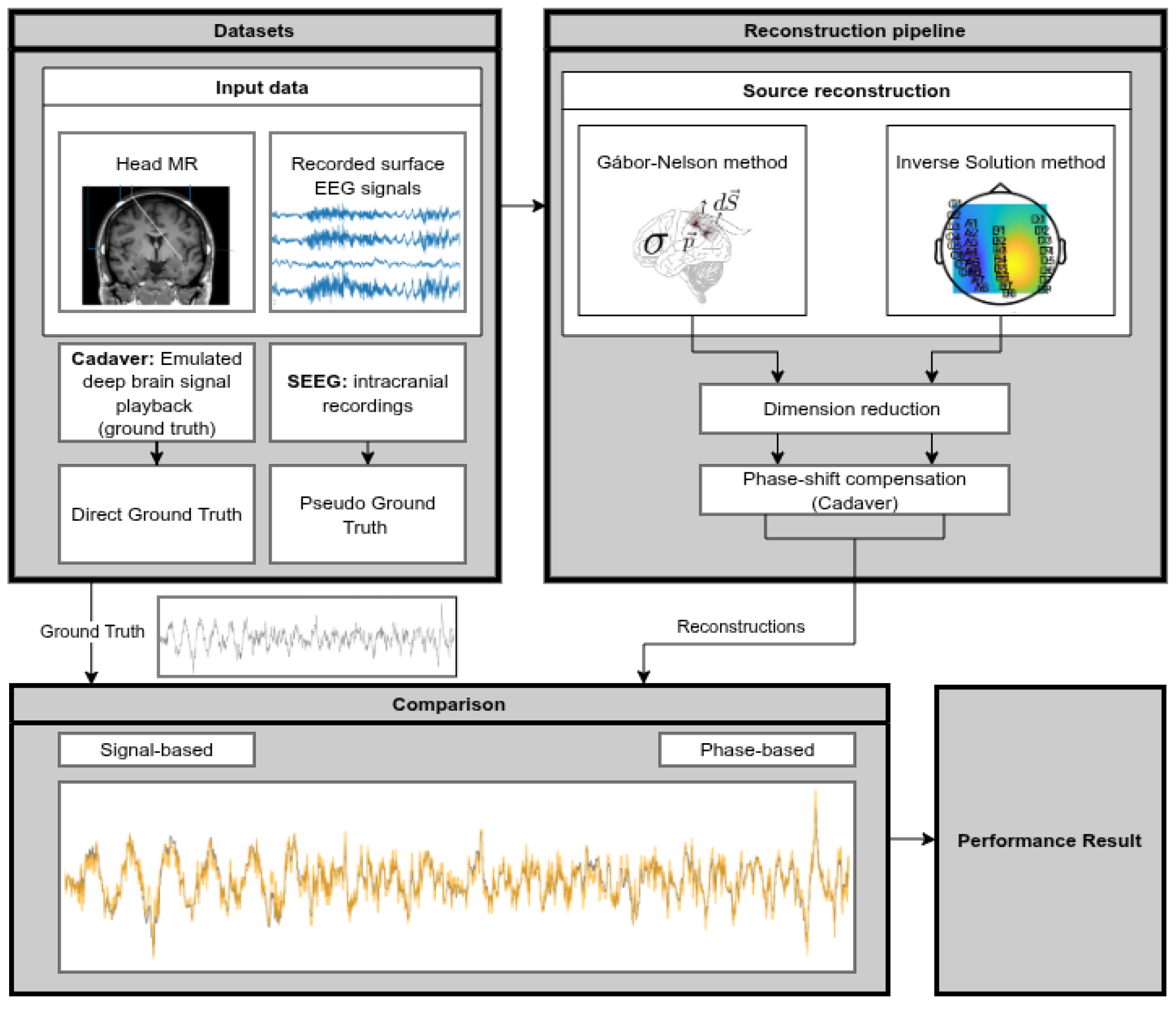
Overview of the phase reconstruction pipeline and validation strategy. The pipeline reconstructs deep-brain phase dynamics from surface EEG using either the Gábor-Nelson or Inverse Solution method. Dimensionality reduction is applied to the 3D dipole time series, followed—in cadaver data only—by frequency-dependent phase-shift correction. Validation is performed using two types of datasets: cadaver recordings with known replayed signals (direct ground truth), and SEEG recordings with pseudo ground truth derived from intracranial electrodes. Final reconstructions are assessed via signal-based and phase-based comparisons to the corresponding reference, yielding performance metrics.

### 2.2 Signal Reconstruction from Surface Potentials

Signal reconstruction was done by applying the Equivalent Current Dipole (ECD) model on the surface potentials. Two methods were employed as dipole modeling: apart from the novel GN method, IS was also applied. Initial n-dimensional EEG recordings are transformed into a Euclidean 3D space [29].

#### 2.2.1 Gábor-Nelson Method

The Gábor-Nelson (GN) method is derived from Gauss’s law and utilizes the surface integration of electric potentials to estimate the net dipole vector within a closed (or approximately closed) measurement surface [24,25]. The method requires knowledge of electrode positions and their surface normals, which were computed from either anatomical meshes or electrode strip geometry.

For a closed surface *S*, the dipole moment 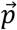 at time *t* can be estimated as based on the measured potential 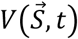 and conductance *σ* as

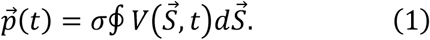

The mathematical background and the experimental application of the GN method are shown in Figure 2.

**Figure 2.**
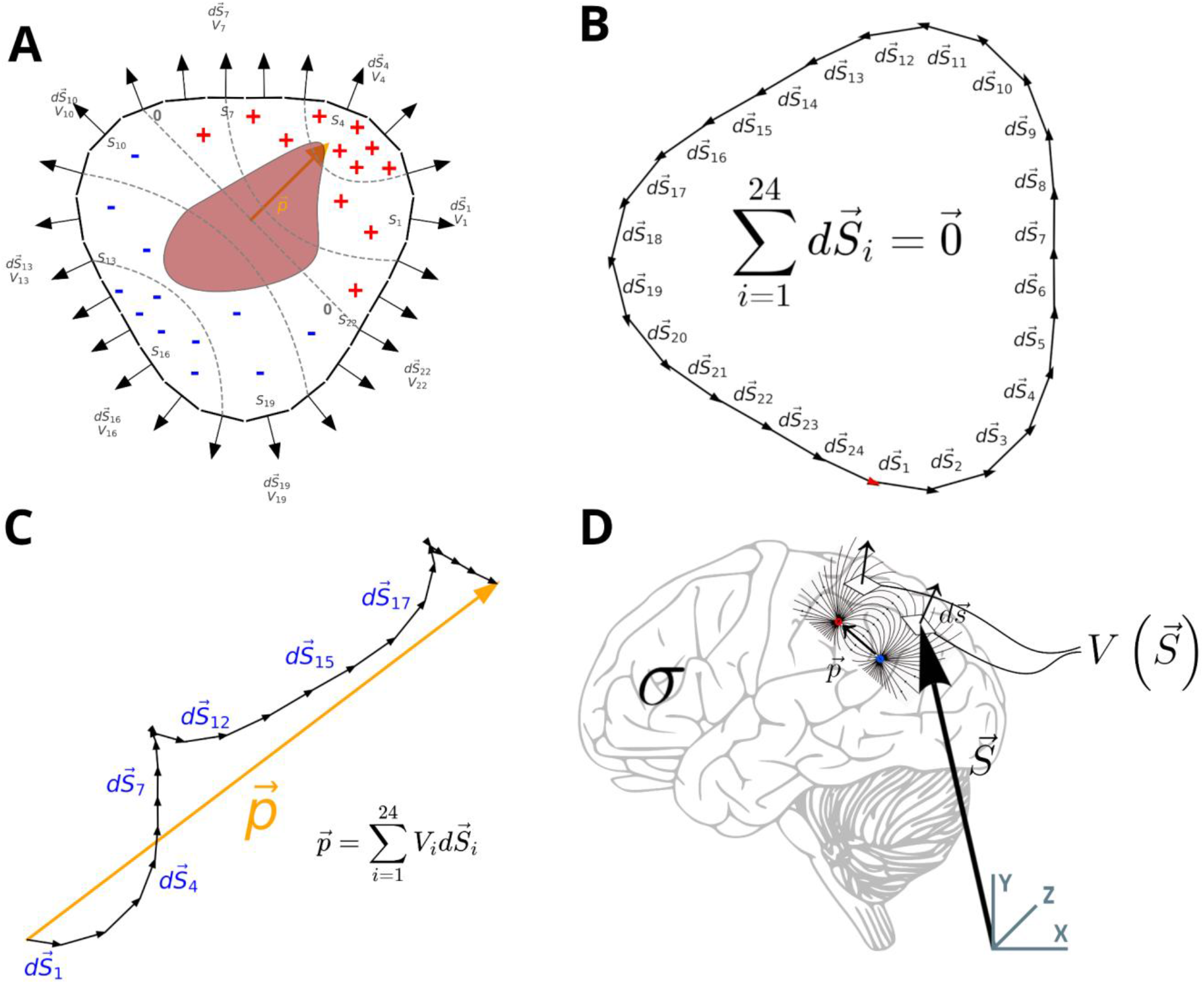
Gábor-Nelson method. A: surface normal vectors positioned around the source. Panel A shows how an enclosed volume’s surface elements (S_1_ … S_12_) and normal vectors 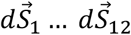 counterclockwise) with their measured potentials (V_1_ … V_12_) are positioned relative to the inner dipole 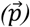. Inner dipoles generate an electric field that is more positive to the right (indicated by the positive signs) and more negative to the left (blue negative signs). The number of signs is correlated with the magnitude of the potential; i.e., the horizontal extremes are the strongest in terms of potential. Furthermore, the resultant vector is shown with a yellow arrow and marked as 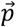, and isopotential surfaces are noted with a dashed line. B: the sum of the individual normal vectors is zero. Panel B indicates that the sum of the individual surface normal vectors is zero if they are unit length. C: the linear combination of the normal vectors, weighted by the local potential, sums up to the resultant dipole vector. It shows that if they are multiplied with the corresponding potential, the sum is a non-zero vector, namely the resultant 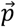 dipole vector. D: Experimental setup. Please note that electrodes were placed on the skull, thus the figure is only for illustration purposes.

In practice, GN was implemented as a discrete summation over the electrodes. Although the theoretical requirement for a fully enclosing electrode array was not met in practice, the residual bias was substantially reduced by subtracting the mean dipole vector over time. The non-enclosing geometry produces a constant offset term given fixed electrode geometry; mean subtraction removes that stationary bias and preserves time-varying dynamics.

#### 2.2.2 Inverse solution with fixed position dipole model

To establish a physically grounded, anatomically informed baseline for evaluating the GN method, a fixed-position inverse solution (IS) was selected as our benchmark methodology for reconstructing dipole source kinetics from multi-channel surface EEG signals [30]. This conventional IS paradigm represents a near-optimal reference standard for equivalent current dipole reconstruction because it systematically integrates subject-specific MRI anatomical data, co-registered electrode configurations, and realistic head volume conductor models to solve the coupled forward and inverse problems of bioelectric field modeling. Furthermore, the algorithm can directly leverage predefined source locations when a priori structural or depth localization metrics are available [31,32]. To ensure mathematically equitable and direct comparison with the lightweight GN estimator, the IS pipeline was explicitly constrained to reconstruct a single equivalent current dipole source. Consequently, by design, this benchmark implementation was restricted from resolving complex, multi-focal or spatially distributed co-activating neural generator

Most common methods for EEG source reconstruction comprises two main components: 1) forward modeling, which involves predicting surface potentials from theoretical source dipole-vectors using a biophysical model, and 2) inverse modeling, which entails estimating the unknown sources responsible for observed potentials. Similarly, the fixed-position IS method begins with forward modeling to produce the leadfield matrix: a matrix of transfer functions that quantifies how dipole orientations contribute to voltage measurements at each electrode. Specifically, for a fixed dipole location inside the brain, the leadfield matrix *L* ∈ *R*^*Nx3*^ defines the mapping from the dipole with normal components in the 3D space to electrode potentials at each of the *N* electrodes. The computed leadfields for each dipole component can be represented graphically, as shown in the topographic heatmaps in Figure 3.

**Figure 3.**
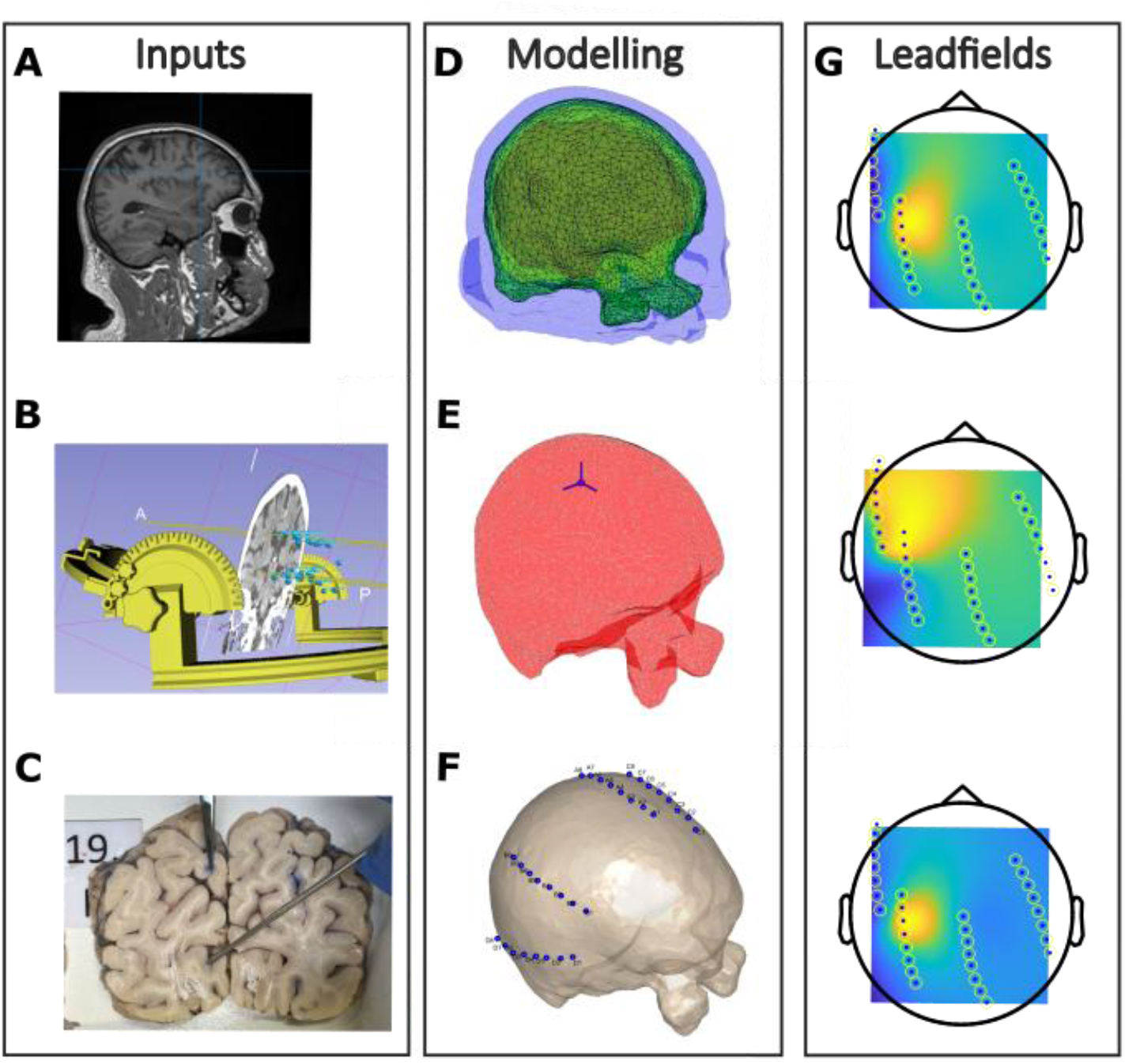
Key elements of the inverse solution method in the cadaver experiment. **A**: Structural MRI of the cadaver head. **B**: Pre-operational planning of the DBS lead trajectories using a custom-built stereotactic frame. **C**: Post-hoc dissection of the cadaver brain, showing the track of the deep electrode penetration. The signal was played as inserted dipole currents on the contact points near the tip of the deep electrode, while the played signal was recorded on a 32 channel electrode system, placed on the surface of the skull and visible on subfigure F and G. **D:** 3D surface meshes representing the different tissue categories of the head based on the T1 and T2 MRI recordings. **E**: Placement of the probe dipole to the location of the signal generator deep-brain electrode. **F**: Placement of the recording surface electrodes. **G**: Modelled leadfields: potentials color coded on the skull surface for the three perpendicular unit dipole directions placed at the known position of the signal-playing electrode. In practice, this illustrates the potential magnitudes at the electrodes generated by three orthonormal probe dipoles with origo shown on panel E.

Assuming a single dipole source and linearity of electromagnetic propagation, the EEG voltage vector *v*(*t*) ∈ *R*^*N*^ at time *t* can be expressed as

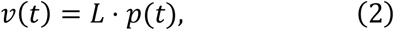

where *p*(*t*) ∈ *R*^3^ is the magnitude of the instantaneous dipole moment vector of a single three-dimensional source. Solving for *p*(*t*) gives:

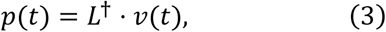

where *L*^†^ is the pseudo inverse of the matrix *L*. Calculating *p*(*t*) for every *k* time instance of an EEG recording *v*(*t* = 1: *k*) predicts the dipole activity in the source and allows obtaining the phase of the source activity from the 3D dipole trajectory that evolves over time.

##### Head Model Construction and Forward Solution

To accurately compute the leadfield matrix *L*, a realistic volume conductor model of the head is required. This model accounts for the layered structure and varying conductivity of the brain, cerebrospinal fluid (CSF), skull, and scalp. We employed the Boundary Element Method [31,33] (BEM) to solve the forward problem due to its computational efficiency and sufficient precision for isotropic, homogeneous compartments.

Computational modeling was performed using customized scripts implemented in MATLAB (The Mathworks, Inc.) and the FieldTrip [34] MATLAB package. The modeling pipeline consisted of the following steps:

1. **MRI preprocessing**: Structural MR images were realigned, resliced, and smoothed.
2. **Segmentation**: Tissue classes (scalp, skull, CSF, brain) were segmented using Statistical Parametric Mapping 12 (SPM12, Wellcome Trust Centre for Neuroimaging, London, UK)
3. **Mesh generation**: Anatomically realistic surface triangulations were computed using the Iso2Mesh algorithm (FieldTrip) or the Neuroelectromagnetic Forward Head Modeling Toolbox [35] (NFT). Parameter tuning, alongside additional surface pruning and refinement, was performed based on visual inspection of the fitted meshes.
4. **BEM model creation**: Using OpenMEEG [36], a four-compartment BEM was constructed, with default conductivities assigned to each compartment.
5. **Sensor and dipole localization**: The locations of the surface (EEG) electrodes were mapped onto the skull (for Cadavers) or onto the scalp (for SEEG subject) mesh, while the location of the dipole source was registered inside the innermost compartment (brain) for both cases. The co-registration of sensor and dipole coordinates with the head model was achieved by applying an affine transformation, aligning both to the MNI coordinate space.

In this study, we modeled the source as a fixed-location dipole with variable orientation and amplitude over time. After calculating the leadfield matrix using the Boundary Element Method (BEM) and re-referencing the leadfield matrix to the EEG reference electrode, the inverse problem was solved to reconstruct the source activity at a fixed-position dipolar source at any given time point. The resulting dipole time series *p*(*t*) is a 3D vector representing momentary source orientation and strength. The temporal evolution of the dipole vector forms a point cloud in three-dimensional space (see Supplementary Figure 5), indicating that the estimated dipole slightly changes orientation over time with additional variability introduced by measurement noise, inaccuracies of the head model, and numerical errors in the forward solution. Consequently, simple vectorial summation of dipole components is insufficient to reliably estimate the true dipole vector. Instead, the dipole orientation and amplitude were approximated by identifying the dominant component within the manifold spanned by the three probe dipole components. The dominant component was computed by applying Independent Component Analysis (ICA) on *p*(*t* = 1: *k*) and subsequently selecting the component exhibiting the highest entropy. We propose that, under high signal-to-noise ratio conditions, dipolar activity is effectively isolated by the independent component aligned with the direction of maximal entropy (Supplementary Figure 4, 5).

##### Validation on Simulated Data

To validate the IS pipeline, a test simulation was constructed, where both the forward and inverse modelling was performed in sequence, to prove that the IS pipeline is capable to reestimate the original input signal of the pipeline with high accuracy. A synthetic (sinusoidal) dipole signal, serving as the ground truth, was projected into three-dimensional space using arbitrary weightings of 5, 10, and 1 for the first, second, and third axes, respectively, to simulate an arbitrary dipole orientation. The projected dipole activity was then forward-modeled using leadfields computed from the head model of a cadaver subject to model the surface signals generated by this synthetic intracranial ground truth source. Gaussian noise was added to simulate measurement noise, see Supp Figure S1, Panel A.

Reconstruction of the intracranial source dipole vector from the simulated surface signals, using the combination of IS and ICA, accurately recovered the original dipole component weightings (i.e., 5:10:1 magnitude ratio). Moreover, the reconstruction yielded a Pearson correlation of r = 0.9999 between the highest entropy independent component and the original ground truth signal. However, there was a 180° phase flip in the ICA-derived signal, due to ICA sign indeterminacy (see Supp Figure S1, Panel B.).

### 2.3 Dimension reduction

The reconstructed dipole signals were inherently three-dimensional. To convert this into a unidimensional signal suitable for phase extraction, Independent Component Analysis (ICA) was applied across the three orthogonal components. For cadaver datasets, a single independent component was extracted, whereas two components were retained for the human clinical dataset. Component selection was based on maximizing signal entropy (Shannon entropy of the amplitude distribution), which empirically favored components with rich temporal structure over background or noise-dominated components. The resulting scalar time series was then used in downstream phase analysis.

A single-component ICA decomposition was applied to cadaver datasets, as the recorded signals originated from a single controlled source (i.e., from an external signal player). For SEEG recordings, a two-component ICA model was adopted to better separate the dominant seizure activity from the background.

To improve model prediction, we used temporal windowing to exclude muscle artifacts, spectral filtering to exclude high-frequency noise and baseline drift.

### 2.4 Phase Correction

In both experimental and clinical EEG systems, the measured signal is subject to frequency-dependent distortions due to the physical properties of the hardware and the possible low-pass filtering imposed by biological tissue [37]. These distortions include amplitude attenuation, which affects signal magnitude and phase delay, which shifts the timing of the signal and is especially detrimental to phase-locked stimulation paradigms.

While amplitude distortion is often less critical for phase-based neuromodulation, phase delay can undermine stimulation timing if not properly corrected. To address this, we implemented a phase correction pipeline that is mapping the frequency dependent phase-transfer function (Bode function) of the experimental setup based on the comparison of replayed and measured signals in the cadaver experiments. Then, we eliminated the hardware-based phase distortion of the signals by fitting the frequency-dependent phase shifts in the Fourier-space.

To correct the phase-delays caused by the recording hardware, a phase-delay function was needed. The phase-delay function, determining the introduced phase-shift with respect to frequency, was calculated in the Fourier space as a difference between the reconstruction and the ground truth as follows.

The reconstructed signal (either by GN or IS) *u*_*r*_(*t*) is transformed into its spectral representation using Fourier transformation

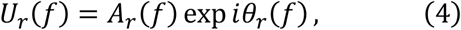

where *A*_*r*_(*f*) and *θ*_*r*_(*f*) are the magnitude and phase at frequency *f*, respectively. The same is applied to the ground truth (GT)

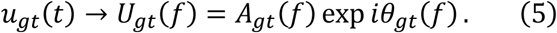

Then the phase difference between the two signals Δ*θ*(*f*) at each frequency is (using circular subtraction)

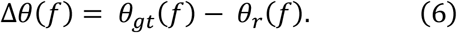

To create continuous phase-delay function, multiple reconstruction-ground truth phase differences were aggregated, and a locally weighted scatterplot smoothing (LOWESS [38]) approximation was applied to all differences to obtain a continuous, general phase-delay function: Δ*θ*_*g*_(*f*). LOWESS is a non-parametric regression method that fits simple models to localized subsets of data to create a smooth curve through noisy observations. At its core, LOWESS repeatedly fits weighted local linear regressions—where nearby points receive higher weights via a kernel function—to estimate a smooth curve that adapts to the data’s local structure. The evaluation is done using leave-one-reconstruction-out cross-validation (LORO-CV). The total recording was segmented by the distinct signal types. To avoid fitting and testing on the same data, when creating a phase-delay function, we omitted the signal segment which was later phase-corrected. No cross-patient data was used, neither were GN and IS reconstructions mixed. For one particular signal segment reconstruction (labelled as 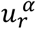, where *α* denotes one segment that is either GN or IS reconstructed), all other segments in that patient and at that location were used to get the phase-delay function 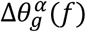. The phase-corrected signal for reconstruction *α* is obtained by adding together the original phases and the phase-correction values at every frequency point as in

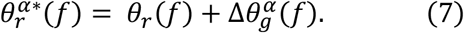

Using inverse Fourier transform with the corrected phases yields the phase-corrected signal 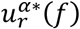. Evaluation was based on the Pearson correlation of the ground truth and the phase-corrected signal 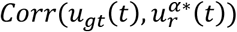. Since the analyzed recordings span less than two minutes, temporal variability of the underlying process is sufficiently limited such that the stationarity assumption should not introduce significant errors.

This method enables phase correction on any complex waveform, regardless of its frequency content, and can correct phase delay in a global and broadband manner.

Fourier-space correction was used to pre-process reconstructed dipole signals prior to phase estimation for cadavers. All residual hardware-related and biophysical phase shifts were compensated for. This correction pipeline was validated across cadaver replay data. SEEG datasets showed no significant delay in phase (see supplementary figure 6), since the recording apparatus was the same for both the surface and the intracranial electrodes. Our correction captures the combined phase response of the acquisition chain under the experimental configuration (electrode interface + amplifier), rather than separating individual contributions.

Furthermore, due to the longer signals being played for cadavers, the signal player’s errors also influenced the measurement. The error in the nominal playback frequency of the device (44.1 kHz) caused the actual frequency to be slightly less than 44.1 kHz. This caused a cumulating lag which only made significant delays for longer measurements for cadavers. Manual correction by resampling was applied.

### 2.5 Evaluation

Evaluation of reconstruction accuracy was done in two distinct approaches: signal-based and phase-based evaluation. Signal-based evaluation consisted of Pearson-correlation between the data points, whilst phase-based evaluation was measured as the mean phase difference of the Hilbert transforms. For Hilbert transform, the reconstructed signals were filtered using a bandpass filter to yield a monotonous time course of phases. The filter limits were established as a 7 Hz wide band centered around the primary frequency component during the statistical testing of the cadaver phase evaluation.

The Hilbert transformation provides a widely used method for calculating the instantaneous phase of a narrowband signal. Given a real-valued time series *u*(*t*), its analytic signal is defined as

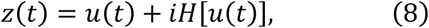

where *H*[*u*(*t*)] is the Hilbert transform of *u*(*t*), which imparts a ±90° phase shift to each frequency component (sign depending on frequency polarity). The instantaneous phase is then

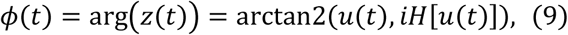

where *ϕ*(*t*) is the instantaneous phase, arg() is the complex argument function and arctan2() is the four-quadrant arcus tangent function. This formulation enables direct tracking of phase progression over time.

In practice, Hilbert-derived phase differences between reconstructed and ground truth signals can be used to estimate **static or dynamic phase delays**. A constant lag between two phase trajectories corresponds to a stable phase delay at a specific frequency, which can then inform correction strategies. Our metric compared the average phase difference between the instantaneous phases of the reconstructed signal and the ground truth.

### 2.6 Experimental Datasets

#### 2.6.1 Cadaver Recordings

The cadaver experiments were approved by the Regional and Institutional Review Board of Human Investigations in University of Szeged (22/2023-SZTE) and the Medical Research Council - Scientific and Research Ethics Committee of Hungary (BM/18167-3/2023). Medical history of cadavers was consulted.

Cadavers were selected based on the absence of major cranial or cerebral abnormalities, confirmed through recent anatomical imaging (CT or MRI), to facilitate accurate modeling of electric current flow. Following prone positioning on the autopsy table, standard cranial measurements were recorded, and electrode entry points were marked using a 10–20 EEG electrode cap (Neuroelectrics NE019-P-NB2). Four, 8-channel electrode strips (Ad-tech TS08R-SP10X-000) were inserted subgaleally through small incisions, secured with sutures, and their position was verified by palpation. A stereotactic frame was then symmetrically affixed to the skull using ear bars. Deep brain stimulation lead placement followed predefined trajectories, planned by co-registering cadaver imaging data with the stereotactic frame model, as detailed in Földi et al. [39]. For source electrode localization, the DBS lead tracks were inked prior to the insertion. The positions of the DBS leads were acquired by tracking back the inked path through the coronal brain slices after post-experimental dissection. These tracks were subsequently reconstructed using pre-mortem MRI in 3D Slicer [40] to obtain the coordinates of the individual current source electrodes.

Cadaver experiments were conducted using three fresh human specimens at the Department of Pathology, Faculty of Medicine, University of Szeged. In each experiment, a modified audio signal player replayed known ground truth (GT) waveforms — composed of repeated seizure fragments — into deep brain tissue via DBS (Deep Brain Stimulator lead, Medtronic 3389) electrodes inserted into multiple locations. Seizure signals were concatenated from multiple recordings acquired during our preclinical study [39]: control seizure periods, along with their adjacent background activity, from four patients diagnosed with focal epilepsy, were chosen as ground truth signals (Table 1). Eleven distinct signal segments were replayed in each cadaver. The signals were delivered at two brain locations in Cadaver 1, three locations in Cadaver 2, and four distinct locations in Cadaver 3. Surface EEG was concurrently recorded via 32 channel electrodes mounted on the skull.

**Table 1.** Dataset summary. Table contains all differences between processed datasets. Cadaver data was tested in three human specimens, whilst SEEG recordings were done in two patients. The number of recordings in total is a magnitude higher for cadavers, just as it is for the total length of recordings (more than an hour for cadavers and only 25 and 20 seconds of seizure activity for SEEG). Phase correction was needed only for the cadaver data. Ground truths were only estimated for the SEEG, and IS was not applicable to Patient B due to missing seizure onset zone location and head model. Evaluated stages consist of before and after phase-correction for cadaver and different frequency bands for the SEEG.

|  | Cadavers | SEEG |  |
| --- | --- | --- | --- |
|  |  | Patient A | Patient B |
| Number of subjects | 3 | 1 | 1 |
| Number of recordings | 11 positions x 9 seizure segments (from 4 patients) = 99 recordings | 1 | 1 |
| Length of the analyzed recordings | ~70 min | 25 sec | 20 sec |
| Phase correction method | Fourier – LOSO-CV | - | - |
| Ground Truth estimation | Replayed previous recordings on intracranial electrode | CSD of SEEG lead ‘F’, penetrating the lesion | Deep dipole estimation with dimension reduction |
| Evaluated reconstructions | GN, IS | GN, IS | GN |
| Evaluated stages | Before-after phase correction | Wide and narrow frequency band | Wide and narrow frequency band |

**Table 2.** Reconstruction comparison. Table contains the comparison of methods listing the benefits and drawbacks of each.

| Factor | Gábor-Nelson | Inverse Solution |
| --- | --- | --- |
| Imaging requirement | Low (optional CT/MR) | High (MRI + segmentation) |
| Tissue categories | Skull curvature (bone/skin) | HD tissue segmentation (bone, skin, WM, GM, CSF) |
| Source localization needed | No | Yes (e.g., lesion or SEEG anchor) |
| Computational burden | Low | High |
| Use case | Acute, fast screening | Planned implantation, surgical prep |

The experimental arrangement using fresh human cadavers helped to eliminate the physiological distortion factors while preserving the physical and biophysical ones, since the biophysical properties (e.g. conductivity, hydration, etc.) and the geometry of the tissues are roughly equivalent to the in vivo circumstances [41], while the active participation of the neuronal networks are not present anymore. Within these experiments the replayed signals were used to test the precision of the reconstruction methods by comparing the reconstructed signal to the ground truth.

#### 2.6.2 Clinical Recordings

We analyzed data from two patients with concurrent SEEG and scalp EEG recordings, both monitored at the University of Pittsburgh Comprehensive Epilepsy Center (UPCEC), Department of Neurology, University of Pittsburgh Medical Center (UPMC; Pittsburgh, PA, USA). The data was acquired in the course of clinical care, and the use for this purpose was approved by the University of Pittsburgh IRB (DUA00004081). All procedures involving human participants were conducted in accordance with the Declaration of Helsinki. The requirement for informed consent was waived by the institutional review board due to the retrospective nature of the study.

Patient A was recorded using 98 contacts SEEG (10 microelectrodes), and 13 scalp EEG electrodes, and patient’s MRI was provided for head model creation and electrode localization. Surrogate clinical data was informing the exact location of the putative seizure-generating lesion, as well as the identification of the SEEG electrode leads position and the contact overlapping with the lesion voxel. Supplementary figure 2 and 3 show a spatial distribution of the available scalp and cranial electrodes. The dataset comprised eight minutes of recording and included a single seizure event lasting 30 seconds. The exact onset and end of the recorded seizure event was annotated and provided by the UPMC medical team.

Patient B was recorded using 87 SEEG contacts and 14 scalp EEG electrodes. Post-operative MRI was provided, but in case of Patient B, the location of the seizure-generating zone was unknown. The provided recording section was 6 minutes long and included a single seizure event of 28 seconds. The intracerebral electrodes were localized and reconstructed from the subject’s post-op MRIs. For both patients, we discarded baseline activity and the initial high-frequency onset parts of the seizures to focus only on the synchronized, low-frequency seizure activity, see Table 1 for more details. Supplementary figure 4 shows a spatial distribution of the electrodes.

For both patients, the signal was bandpass filtered between 1–80 Hz using a fourth-order, zero-phase Butterworth digital filter, and a 60 Hz notch filter was applied using a second-order IIR design to filter power-line noise. For the evaluation phases, a wideband 1-30 Hz and a narrowband (seizure frequency dependent) filter of the same kind were also used.

#### 2.6.3 Ground Truth determination

In the lack of ground truth data (i.e., the exact seizure generating dipole), GT was approximated from the SEEG signals in the Clinical Dataset using multiple approaches with different tradeoffs between accuracy and complexity.

In Patient A, as a first approach, we selected the SEEG channel (i.e., channel 5 of the SEEG shank F) with the largest spectral power in the range of epileptic activity (3-20 Hz). Channel 5 of shank F was also depicted by a trained medical personnel as the ‘seizure source’ signal with no prior knowledge of our selection. Channel F5 was also the nearest passing shank to the reconstructed cortical lesion location. The SEEG contacts along shank F were evenly spaced, allowing approximation of the one-dimensional current source density (CSD) via a discrete second spatial derivative. Boundary contacts were excluded from the analysis, and no additional spatial smoothing was applied. Under these assumptions, the second spatial derivative provides a standard estimate of local transmembrane current density along the electrode axis. The second spatial derivative acts as a spatial high-pass filter, attenuating distal changes, such as artifacts arising from distant sources that affect all contacts similarly, and enhances local activity. The reconstructed source signals, approximated using both the Inverse Solution and Gábor–Nelson methods, were evaluated against the two ground truth dipole estimates: the signal from the F5 shank and channel 5 and the one-dimensional CSD. The one-dimensional CSD provided a significantly more accurate approximation of the ground truth and was therefore used as the ground truth signal throughout the analysis.

In Patient B, due to the absence of prior lesion localization and a defined seizure onset zone, a moment-based dipole estimate derived from the intracranial recordings was used as the ground truth. Electrode position vectors were weighted by their corresponding potentials and summed across contacts, yielding a time-varying three-dimensional dipole vector. Principal component analysis was applied to the resulting dipole trajectory, and the dominant component was selected as the ground truth reference. This estimate was further validated by trained medical personnel through comparison with the concurrent surface EEG recordings, confirming that the same seizure-related phenomenon was present in both modalities.

## 3 Results

### 3.1 Cadaver Validation

#### 3.1.1 Signal-based evaluation

We first applied the GN and IS methods to cadaver recordings, which included surface EEG recordings generated by replayed seizure waveforms through deep brain stimulation (DBS) electrodes (Figure 5B). This dataset was selected for its anatomical fidelity, replay consistency, and compatibility with both reconstruction methods. Phase delay correction was used in the 1-100 Hz regime (Figure 5C).

**Figure 5.**
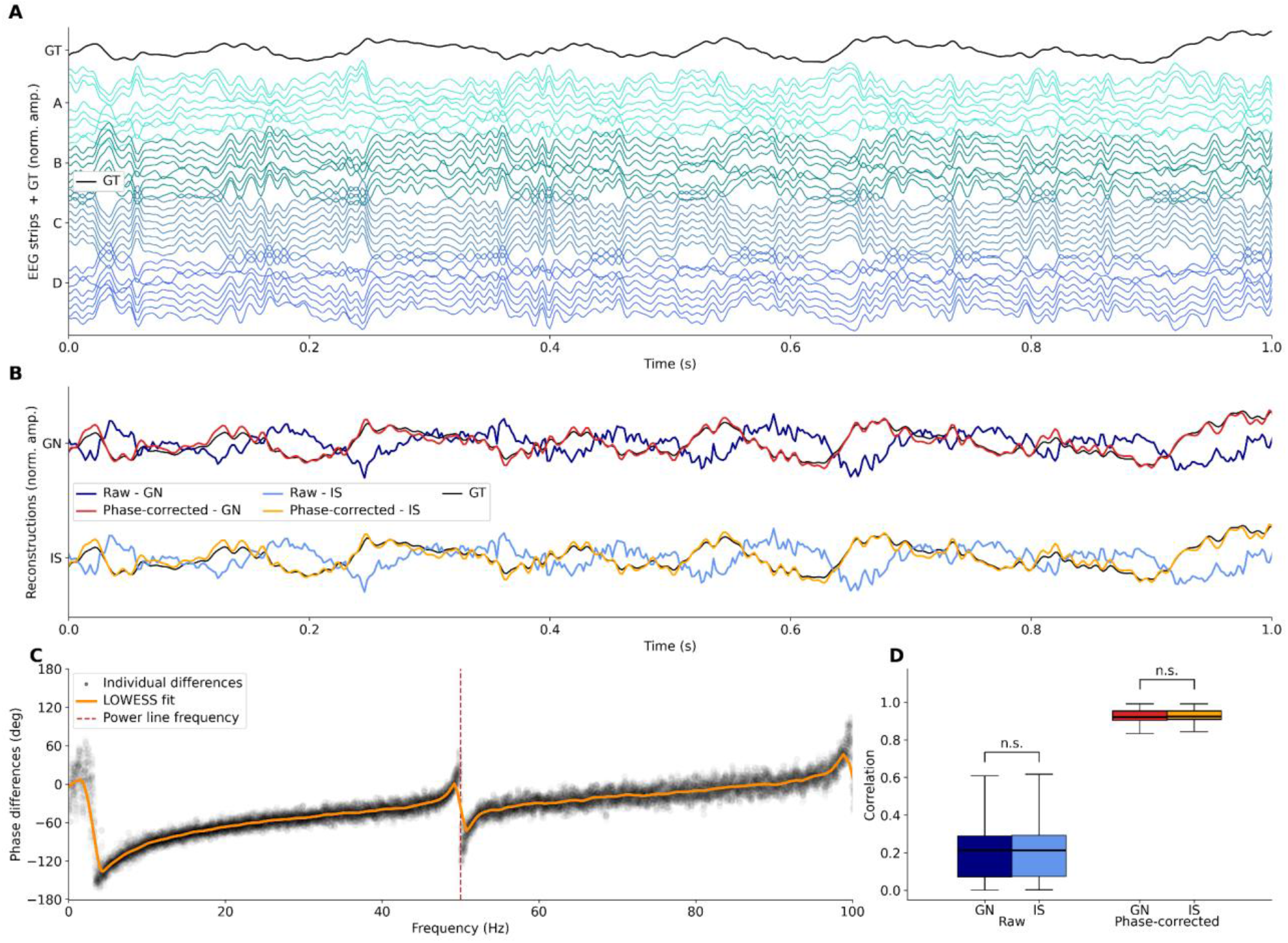
Signal reconstruction. **A.** Recorded voltages per strip and Ground Truth signal. The 32 EEG channels are grouped to strips A, B, C and D are plotted in separate colors; each strip shows 8 channels. The first row shows the ground truth (GT) source in black. **B.** Example signal played and reconstructed via the Gábor-Nelson and Inverse Solution methods. Raw reconstructions (dark and light blue) showed limited correlation to the ground truth (GT, black); however, with phase correction, these increased to 0.91 (noted in red and orange). **C.** Example of the phase correction function. Individual Δθ(f) phase differences are marked with dots, whilst the general continuous Δθ_g_(f)fit is marked with blue. **D.** Phase correction yields a large, method-agnostic improvement in reconstruction consistency. Boxplots of voxel-wise correlation coefficients between repeated reconstructions under Raw (no phase compensation) and PhCorr (with phase correction/compensation) conditions for the Gábor–Nelson (GN) and inverse-solution (IS) dipole-modeling methods. In the Raw state, both GN and IS show low median correlations (~0.21), which rise to high consistency (~0.91) after phase correction. Paired Wilcoxon signed-rank tests confirm that phase correction produces a highly significant boost in consistency (Raw vs PhCorr, p < 0.001), while paired comparisons of GN vs IS in either Raw or PhCorr regimes are non-significant (n.s., p > 0.05), demonstrating that both modeling methods benefit equally from phase correction

In total, 99 recording segments were taken into the analysis, as detailed in section 2.6.1. Dipole reconstruction was done using the GN and the IS methods separately on each recording. Before phase-correction, both GN and IS methods yielded an average correlation of r=0.212. After phase correction, the average correlation increased substantially to r=0.911 and r=0.913 for the GN and IS methods, respectively (Figure 5D). The reconstruction accuracy of the two methods did not differ significantly at either reconstruction stage (paired Wilcoxon signed-rank test, p>0.05 in both cases).

#### 3.1.2 Phase-based evaluation

Momentary phase agreement between ground truth and reconstructed signals was also quantified. An example of the corrected phases is shown in fig 6A. PhCorr corrected the bimodal polar distribution of the raw phase differences into one distribution, centered around 0 (Fig 6B). Raw errors during the unit-based phase comparison resulted in a distribution with a mean around 90°, while PhCorr errors form a narrow, symmetric peak at 0° (red dashed line) – regardless of reconstruction method. The polar means and the dispersions of the phase differences of the reconstructions from GT decreased significantly after PhCorr (Fig 6B). There was no significant difference between the reconstruction methods (i.e., GN vs IS). Aggregated mean ± SEM phase offsets (Raw: GN = 90.6 ± 53.7°, IS = 91.7 ± 53.3°; PhCorr: GN = 5.1 ± 33.1°, IS = 6.1 ± 32.3°) demonstrate that phase correction significantly reduces both bias and variability.

**Figure 6.**
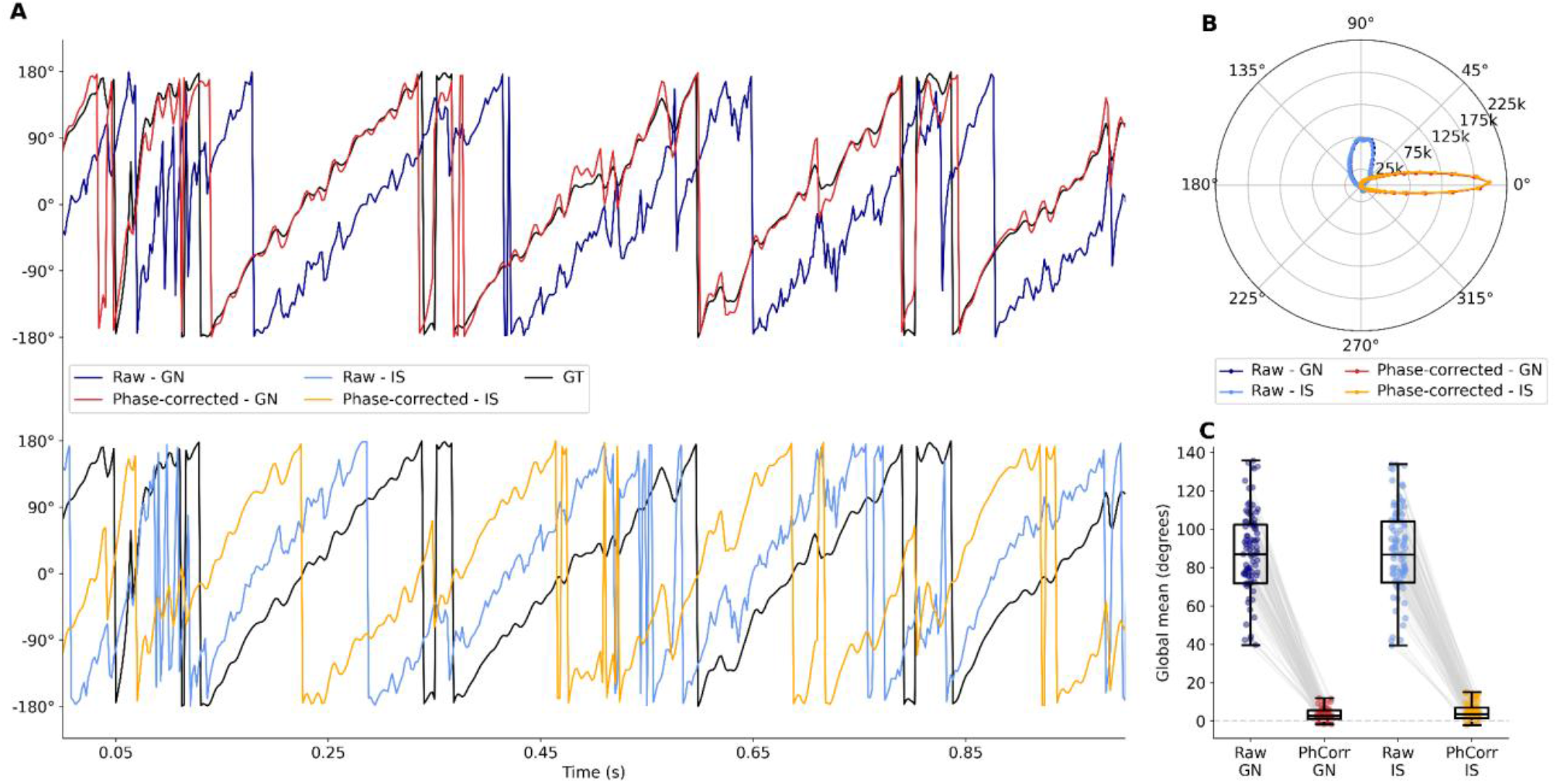
Phase analysis of the Cadaver dataset. **A.** Example of reconstructed signal phases for the Gábor-Nelson and Inverse Solution methods. Raw reconstructions (dark and light blue) showed significant average phase difference (around −70°) to the ground truth (GT, black), however, with phase correction, these decreased to −1.38° and −0.44° (noted in red and orange). **B.** Rose plots of phase-difference histograms show that, in the Raw condition, both Gábor– Nelson (GN) and inverse-solution (IS) reconstructions exhibit broad phase distributions with a mean of ~90°, whereas phase correction (PhCorr) collapses errors into a tight cluster around 0°, with means around −8°. **C.** Cycle-based phase error between the reconstructions and the GT. The central line represents the median; the box spans the interquartile range (IQR), and the whiskers extend to 1.5×IQR. Outliers are not shown. Data points of raw and phase corrected pairs are connected with grey lines.

Statistical significance testing on the phase differences was performed on unit-level phase differences between each reconstruction and the ground truth (GT). Both the GT and reconstructed signals were band-pass filtered using a 7 Hz wide bandwidth filter, centered on the dominant frequency component, and segmented into units corresponding to individual oscillatory cycles based on the first peak of the GT autocorrelation function. For each unit, the mean phase difference between the reconstruction and the GT was computed, yielding 154–600 units per signal (dominant frequencies spanned 5-12 Hz). Differences between reconstructions were assessed using the Mardia–Watson–Wheeler test to determine whether unit-level phase differences originated from the same circular distribution. No significant differences were observed between the GN and IS methods at either the original or phase-corrected stage across any of the 99 signals (all p > 0.1). In contrast, phase correction produced significant changes in the distribution of unit-level phase differences for both GN and IS reconstructions (all p < 0.001). This procedure provides a robust comparison by accounting for temporal autocorrelation, circularity of phase data, and distribution-level differences without parametric assumptions. Global mean errors between reconstructions and the ground truth are shown in figure 6C (Raw: GN = –74.3 ± 50.2°, IS = –75.0 ± 50.2°; PhCorr: GN = –4.3 ± 4.8°, IS = –5.0 ± 5.1°).

### 3.2 Validation on clinical dataset

Validation on the clinical SEEG dataset was also conducted in two ways: signal-based correlations and phase-based average difference with Hilbert transform.

#### 3.2.1 Patient A

For the Inverse Solution, a full 4-layer BEM model was generated from the patient’s MRI, and the IS was employed to the scalp signal. The true dipole was identified as the highest entropy ICA component for both the GN and the IS method. Since both the deep and surface electrodes were recorded with the same hardware, introducing the same phase shifting to both signal types, there was no system-dependent net phase lag between the SEEG and scalp EEG signals. Two reconstruction types were assessed: one with a wide ranged frequency band (1-30 Hz) and a narrower (12-18 Hz) tuned to the dominant seizure oscillation.

Scalp electrode selection was based on anatomical geometry, volume-conduction properties of deep sources, and artifact susceptibility. Based on a priori information provided by the clinicians, the epileptogenic region was located deep in the right posterior–inferior brain, for which central and midline electrodes provide best sensitivity from the available ones to seizure-related synchrony. The signals of *C3, C4, Cz, Fz, Pz* electrodes were used in this analysis. These electrodes are also less affected by facial muscle artifacts compared with frontal and inferior frontal–temporal sites. Frontal pole electrodes were excluded due to blink- and drift-related contamination, while lateral frontal–temporal electrodes were omitted because of frequent EMG dominance and limited added information for deep posterior sources. This selection reduces artifact-driven variance while preserving robust sensitivity to seizure propagation.

The annotated seizure starts with high frequency, low amplitude oscillation which is not detectable using scalp electrodes with the electrode topography of the current measurement. Accordingly, the analyzed signal was truncated to a 25-second segment characterized by high-amplitude, low-frequency synchronized activity (Supplementary figure 7).

Signal-based comparisons (figure 9A-B) showed high correlation in both GN and IS cases for every frequency setup. For the wider frequency spectrum, GN and IS yielded r=0.632 and r=0.643 correlations, whilst for the narrower, both increased to r=0.8, indicating that the same neural activity has been reconstructed, but with slightly different background noise. Phase-based analysis indicates that restricting the frequency range increases the mean phase offset relative to the GT while substantially reducing dispersion for both reconstruction methods. Aggregated circular mean ± dispersion values were: Wide—GN: 16.8 ± 60.7°, IS: 18.3 ± 59.6°; Narrow— GN: 27.6 ± 33.1°, IS: 28.0 ± 33.1°. Mardia–Watson– Wheeler tests revealed no significant difference between GN and IS neither in the wideband condition (p = 0.993), nor in the narrowband condition (p = 0.994) (Fig. 9C–D).

**Figure 8.**
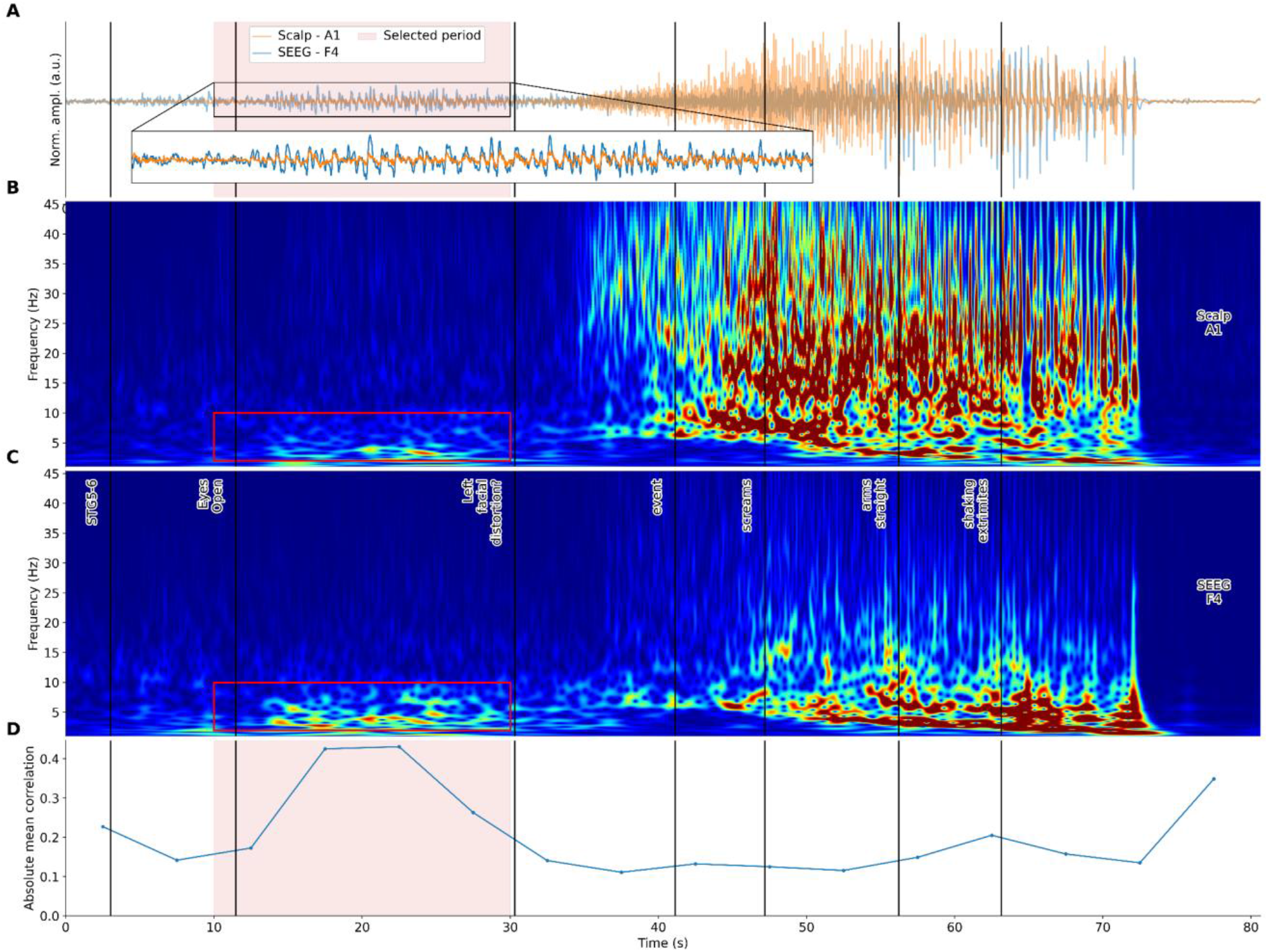
SEEG Patient B. **A.** EEG of the selected channels from the scalp (A1) and SEEG (F1). **B-C.** Wavelet transformations of the whole annotated seizure. The two subplots show wavelet transformations for the highest-amplitude electrodes, one scalp (A) and one SEEG (B). The color scale highlights the 10–30 second region in both cases. Additional annotations from the EDF files are marked with black vertical lines. Higher amplitudes are represented in red. The chosen period and narrower frequency regime are highlighted with a red bounding box. **D.** Pearsons’ correlation coefficient average values between the deep and surface channels, comparing all-to-all. The comparison was performed in 5 seconds-long windows. Prior to calculating the mean, the absolute values of the correlation coefficients were calculated. The selected period between 10-30 seconds is highlighted in red.

**Figure 9.**
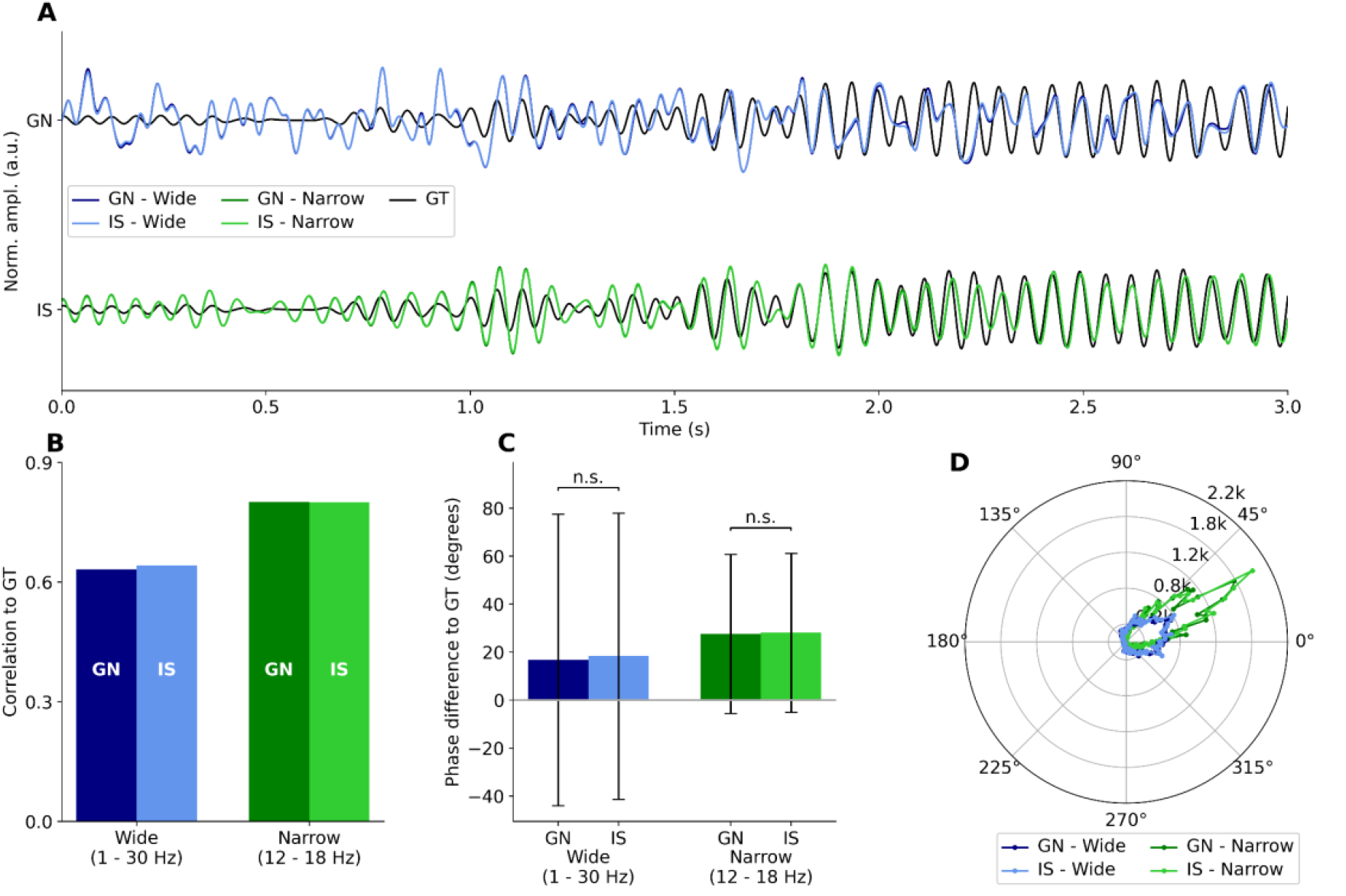
SEEG Patient A summary. **A.** The two reconstruction methods were also assessed under a wider (1-30 Hz) and a narrower (12-18 Hz) frequency range. The beginnings of the signals by Gábor-Nelson method are shown with darker shade and the Inverse Solution with lighter colors (marked as GN and IS, respectively). **B:** Bar graph of the total correlation between the reconstructions and the GT. **C.** Corresponding aggregated circular mean and dispersion values; a significant GN–IS difference is present only in the wideband condition (Mardia–Watson-Wheeler test). **D.** Rose plots illustrating phase-difference distributions for Gábor-Nelson (GN) and Inverse Solution (IS) reconstructions under wide and narrow frequency conditions, showing reduced circular dispersion and a higher mean phase offset in the narrowband regime.

#### 3.2.2 Patient B

Dipole reconstruction from the measurement of Patient B was performed in the same manner as for Patient A, except that only GN modeling was employed due to the absence of clinical information regarding the seizure onset zone, which is essential for applying the IS method. Similarly to Patient A, the recording was truncated to a 20-second segment exhibiting high amplitude oscillation and elevated spectral power within the seizure frequency range (1– 30 Hz).

Temporal and spectral filtering, as well as the selection of the analyzed 20-s segment, were guided by clinician annotations and prior inspection of the time–frequency structure of the recording (Supplementary Fig. 8).

Dipole reconstruction using Gábor–Nelson modeling combined with ICA-based dimensionality reduction demonstrated a high correlation with the estimated ground truth. The strongest correlations between EEG channels and the GN reconstruction were observed at channel P9, T5, and P3, suggesting more widespread cortical activity. (Figure 10A). Signal-based comparisons showed that the wideband reconstruction (1–30 Hz) yielded a correlation of r=0.720, whereas restricting the frequency range to 2–10 Hz resulted in a slightly higher correlation of r=0.758, indicating that the broader frequency range contains marginally higher noise relative to the underlying neural activity (fig 10B). Phase-based analysis revealed comparable phase-difference distributions across frequency bands, with aggregated circular mean ± dispersion values of −9.9 ± 60.7° for the wideband condition and −12.6 ± 57.4° for the narrowband condition, indicating a modest increase in mean phase offset and a reduction in dispersion under the narrower frequency regime (fig 10C-D).

**Figure 10.**
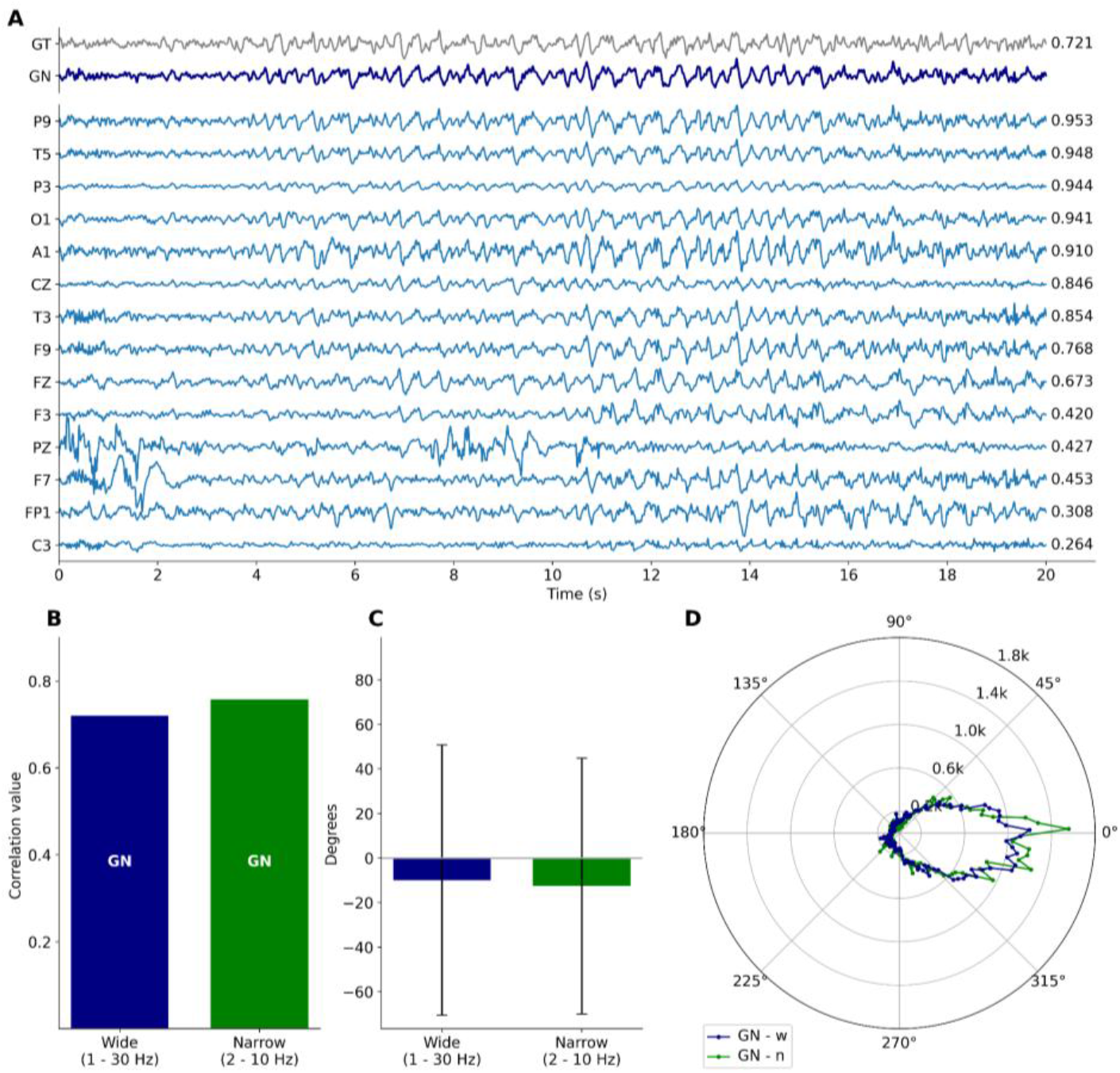
SEEG Patient B summary. **A.** Surface EEG signals (14 channels) shown alongside the deep dipole ground truth (GT, gray) and the Gábor–Nelson (GN, dark blue) reconstruction; correlation values relative to GN are indicated. **B.** Signal-based comparison of GN reconstructions under wide (1–30 Hz) and narrow (2–10 Hz) frequency ranges, showing only a modest increase in correlation with frequency restriction. **C.** Aggregated circular mean and dispersion phase offsets, illustrating a slight increase in mean phase difference and reduced dispersion for the narrowband condition. **D.** Rose plots of phase-difference distributions demonstrating similar phase structures across frequency bands.

## 4 Discussion

### 4.1 Summary of findings

This study presents a comprehensive methodology for reconstructing the phase of deep epileptic oscillations from non-invasive EEG signals. We validated the framework across two fundamentally different domains: controlled cadaver experiments and in vivo SEEG and concurrent scalp EEG recordings from epileptic patients. Our pipeline— which combines dipole reconstruction, dimensionality reduction, and frequency-dependent phase correction—demonstrated robust signal and phase fidelity in both settings, underscoring its potential clinical applicability in seizure phase monitoring and phase-locked stimulation.

While contemporary literature predominantly concentrates on macro-level tasks of detecting, predicting, or spatially localizing intracranial activity from surface recordings, this study expands the current state of the art by evaluating the temporal fidelity of source-level phase reconstruction. This paradigm shift is vital for clinical closed-loop neuromodulation. Successfully executing phase-locked stimulation fundamentally demands that the underlying oscillatory phase can be recovered with sufficient temporal precision to reliably guide microsecond-level stimulation timing. In this context, our results directly complement and extend the cross-modal validation frameworks established by several pivotal recent studies [7,13,42,43]. By shifting the primary benchmarking target from purely spatial correspondence to temporal phase accuracy, this work addresses a critical, under-explored dimension of non-invasive electrophysiology.

In cadaver experiments, phase correction proved indispensable for biologically meaningful phase estimation. Following the correction of hardware-induced phase delays, both Gábor–Nelson (GN) and inverse-solution (IS) reconstructions achieved high signal similarity to ground truth, with correlations exceeding r=0.91 and mean phase offsets reduced to approximately 8–9°. Importantly, while high correlation indicates strong waveform similarity, it does not imply a small phase difference by itself; the combined improvement in correlation and reduction in phase offset confirms that phase correction effectively aligns the reconstructed signals in both amplitude and timing.

The cadaver replay paradigm overcomes a profound limitation to most human scalp-intracranial validation studies: the inability to isolate an exact, deterministic ground truth signal entirely free from biological background noise. By electrically driving known, pre-recorded seizure waveforms through deep intracerebral electrodes, the surface-reconstructed trajectories could be compared directly with the pristine, uncorrupted deep source activity. This high degree of experimental control enabled us to mathematically quantify not only waveform similarity, but also the absolute phase error remaining after compensating for frequency-dependent hardware delays. Consequently, the cadaver dataset serves as an idealized, highly controlled technical benchmark for phase retrieval under optimal signal conditions. This stands in sharp contrast to clinical simultaneous scalp–SEEG recordings, where the true underlying generator cannot be directly observed and can only ever be approximated from adjacent intracranial contacts.

Validation using human SEEG recordings confirmed the feasibility of the approach under clinically realistic conditions, where true ground truth is not directly accessible. Instead, reference signals were estimated using second spatial derivatives for Patient A and deep dipole modeling for Patient B. Despite these constraints, narrowband reconstructions yielded consistently high performance. For Patient A, both GN and IS methods achieved correlations of approximately r≈0.8 with similar mean phase offsets (~17–18°), indicating that the GN method can achieve accuracy comparable to IS when a reliable reference is available. For Patient B, where precise anatomical localization and seizure onset zone definition were unavailable, GN reconstruction achieved correlations up to r = 0.76 with moderate phase dispersion, demonstrating that meaningful phase tracking remains possible even in cases constrained by the absence of MRI, uncertainty in source localization, or the need for accelerated computation.

In contrast to the idealized conditions of the cadaver assays, the simultaneous clinical scalp–SEEG recordings provide a substantially more realistic and translationally relevant test case. These in vivo datasets are inherently challenged by ongoing background activity, variable patient-specific anatomy, constrained scalp electrode coverage, and pseudo-ground-truth reference that must be mathematically estimated from localized intracranial contacts rather than an uncorrupted, known input waveform. Given these clinical constraints, the reconstruction metrics achieved here should be interpreted as an initial proof-of-principle demonstration rather than definitive clinical validation. Nevertheless, the capacity to reliably recover source-level phase dynamics under these noise contaminated conditions underscores the clinical feasibility of translating this framework beyond controlled replay experiments. These findings motivate future, larger-scale validation spanning a broader diversity of patient populations, distinct seizure types, and varied electrode configurations.

The ability to track deep seizure phase non-invasively opens avenues for phase-locked stimulation with minimal invasiveness along with personalized seizure detection algorithms that exploit dynamic phase features and adaptive neuromodulation protocols guided by real-time EEG phase estimates. Even though this study was conducted offline, online phase tracking is also manageable using a real-time phase-tracking algorithm [44].

In contexts where invasive SEEG implantation is not viable or only temporary, GN-based surface estimation can serve as a longer-term phase-tracking method, and the IS method-based pipeline may further enhance phase estimation accuracy.

Our analysis reveals complementary strengths between the two reconstruction approaches:

We recommend using the IS method when full anatomical models and source priors are available, however, in settings where only surface recordings are available or imaging is missing, GN offers a viable alternative with competitive accuracy, while it does not require source-location priors or MRI-derived leadfields, significantly reducing computational needs for phase-locked stimulation.

### 4.2 The validity of the dipole approximation

The modeling of bioelectric activity in the brain often adopts the Equivalent Current Dipole (ECD) as the canonical source representation [29]. This choice arises from both the physical and physiological properties of extracellular brain fields, and the mathematical formulation of electromagnetic field propagation in biological tissues. According to volume conductor theory, brain electric fields arise from transmembrane current sources. Under biologically relevant frequencies (approximately 1– 100 Hz, encompassing typical EEG and SEEG frequency bands), the quasi-static approximation is valid because displacement currents remain negligible compared to conductive currents in brain tissue. This approximation, combined with the principle of charge conservation, implies that isolated current sources or sinks—current monopoles— cannot exist biologically [45]. Consequently, the first non-vanishing term in the multipole expansion of the electric field generated by neural sources is typically the dipole term [46]. However, if the dipole moment of the source distribution is negligible—i.e., the distribution is highly symmetric—the electric field is determined solely by higher-order terms.

Since the electric field and potential generated by these higher-order terms decay more rapidly with distance than those of a dipole, a localized spatial distribution of neural current sources can generally be well approximated by an equivalent dipole, but only from a sufficiently large distance [26]. If, on the other hand, the spatial extent of the sources is comparable to or larger than the electrode array (e.g., reverberating networks), more complex models— potentially involving multiple dipoles or higher-order terms (e.g., quadrupole, octupole)—may be required to accurately describe the source distribution. Thus, the validity of the dipole approximation strongly depends on the spatial extent and configuration of the current sources, as well as the extent and relative position of the recording electrode system [47–49].

Cosandier-Rimélé et al. [50] have attempted to delineate the minimum cortical surface area required to generate scalp-detectable discharges. These quantitative modeling studies have shown that scalp EEG is preferentially sensitive to spatially extended and temporally coherent cortical source patches, whereas much smaller active areas can be detected intracranially. For comparable spike-to-background ratios, the effective cortical area contributing to scalp-detectable activity increases substantially relative to intracerebral recordings. Importantly, this reflects the extent of synchronized neural activity rather than anatomical lesion size, and from a distance such coherent source distributions increasingly suppress higher-order multipole contributions, reinforcing the dominance of the dipole term in scalp measurements.

Although the single-dipole approximation is adequate for describing the electric potential generated by the heart—and is therefore widely used in the cardiac literature—it is typically insufficient to characterize the complex neural activity of the brain [51]. While this limitation holds true for normal brain function, focal epileptic seizures often originate from a confined cortical region, characterized by highly synchronized firing of aligned pyramidal neurons. Such localized synchronization generates a coherent dipolar current source that can dominate over diffuse or asynchronous background activity during the earliest stages of seizure onset, and this synchronization raises the possibility that the electrical activity of the epileptic zone may be well approximated by a single dipole—at least during the earliest phase of the seizure. If this assumption holds, and the single-dipole model proves to be a valid approximation in most cases and patients, then the dipole parameters could be estimated using simpler or more constrained source localization techniques that require fewer parameters and offer greater robustness.

Moreover, the single-dipole description allows us to define and track a single phase of the epileptic oscillation, which could subsequently be exploited in phase-locked neuromodulation techniques. Importantly, the success of single-dipole modeling in this study does not imply that epileptic networks are intrinsically dipolar, but rather that, from the perspective of scalp measurements, synchronized seizure activity can be well approximated by a dominant dipolar component.

### 4.3 Limitations and Considerations for Future Work

Several limitations of the present study warrant consideration. First, while the cadaver experiments provided an exact ground truth, validation in SEEG recordings necessarily relied on surrogate references derived from spatial derivatives of the measured signals. Although such approaches are well established, more direct biophysical modeling (such as full three-dimensional current source density (CSD) estimation) could further strengthen ground truth approximation in future studies. Second, the use of ICA-based dimensionality reduction introduces an inherent polarity ambiguity, as these methods yield components with arbitrary sign. Although waveform skewness provided a generally reliable criterion for post hoc polarity correction, this approach may fail for highly symmetric or balanced oscillatory activity.

Additional limitations arise from the geometry of SEEG recordings. The irregular spacing and orientation of intracranial electrodes constrain the accuracy of higher-order spatial derivative estimates, such as full 3D Laplacians, particularly in regions near sulci or fluid-filled cavities. Furthermore, while the present pipeline was evaluated in an offline setting, translation to real-time operation would be highly desirable for closed-loop neuromodulation applications. In this context, an important open question concerns the frequency dependence of phase delays. Systematic investigation of phase shifts across a large SEEG–scalp dataset spanning diverse seizure frequencies could inform whether frequency-specific compensation is required. Finally, extending the framework to allow dynamic, time-varying dipole estimation would enable modeling of seizure propagation and spatial evolution, offering a more complete description of epileptic dynamics.

Although single dipole-based methods, like the Gábor-Nelson method are rarely used for EEG source localization, these results showed that the single dipole approximation was good enough to capture the majority of the epileptic deep source activity based on the scalp recordings. Moreover, the Gábor-Nelson method achieved similar precision in the reconstruction of the deep signals and their phases without MRI-based tissue segmentation as the fixed-dipole inverse solution that utilized computationally intensive lead-field calculation on an inhomogeneous head model.

Together, these results lay the foundation for non-invasive seizure phase monitoring, a critical prerequisite for safe and effective phase-locked therapeutic interventions in epilepsy. Sex- and gender-based analyses were not performed due to the limited sample size and the methodological focus of the study, which may limit generalizability.

## CRediT authorship contribution statement

**Kristóf Furuglyás:** Formal analysis, Investigation, Methodology, Software, Visualization, Writing – original draft, Writing – review and editing. **Márton Huszár-Kis:** Formal analysis, Investigation, Methodology, Software, Visualization, Writing – original draft, Writing – review and editing. **Bálint Horváth:** Data curation, Writing – review and editing. **Andrea Pejin:** Data curation, Writing – review and editing. **Nóra Forgó** Data curation, Writing – review and editing. **István Langó:** Resources, Writing – review and editing. **Shobhit Singla:** Data curation, Writing – review and editing. **Márton Görög:** Software, Writing – review and editing. **Péter Vass:** Software, Writing – review and editing. **Zoltán Chadaide:** Data curation, Validation, Writing – review and editing. **Tamás Laszlovszky:** Project administration, Supervision, Writing – review and editing. **Orrin Devinsky:** Conceptualization, Writing – review and editing. **Anto I. BagiĆ:** Data curation, Writing – review and editing. **Zoltán Somogyvári:** Conceptualization, Formal analysis, Investigation, Methodology, Supervision, Writing – original draft, Writing – review and editing. **Antal Berényi:** Conceptualization, Formal analysis, Funding acquisition, Investigation, Methodology, Supervision, Writing – original draft, Writing – review and editing.

## Declaration of generative AI and AI-assisted technologies in the manuscript preparation process

During the preparation of this work, the authors used OpenAI’s ChatGPT for language improvement. After using this tool/service, the authors reviewed and edited the content as needed and took full responsibility for the content of the published article.

## Data availability

Data will be made available upon reasonable request.

## 5 Supplementary Material

### 5.1 Simulation

**Supp Figure 1.**
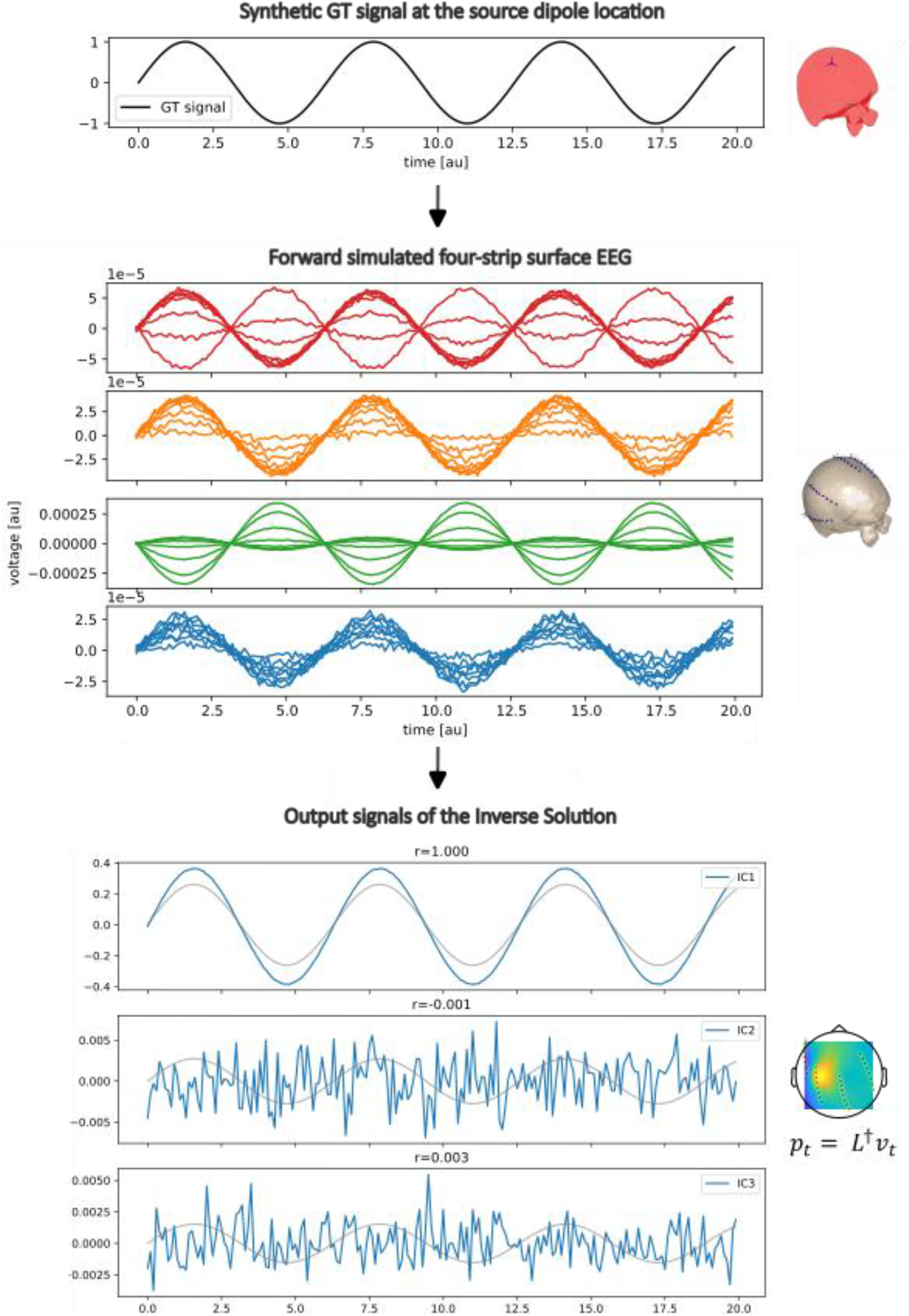
Validation of IS method on simulated dataset A. A sinusoidal ground truth (GT) signal that serves as the source dipole, positioned at a theoretical intracerebral location. **B.** Forward simulation: rows 1-4 illustrate the EEG signal generated from the GT signal using the patient-specific forward model of Patient 1. The 32 EEG channels are grouped into strips A, B, C and D, reflecting the actual electrode layout, and are plotted in separate panels accordingly. **C.** Inverse Solution: Independent component embedding of the three dipoles reconstructed by the Inverse Solution method are shown. Linear correlation coefficients between the Independent Components (ICs) and the GT signal are indicated above each corresponding subpanel.

### 5.2 Datasets

**Supp Figure 2.**
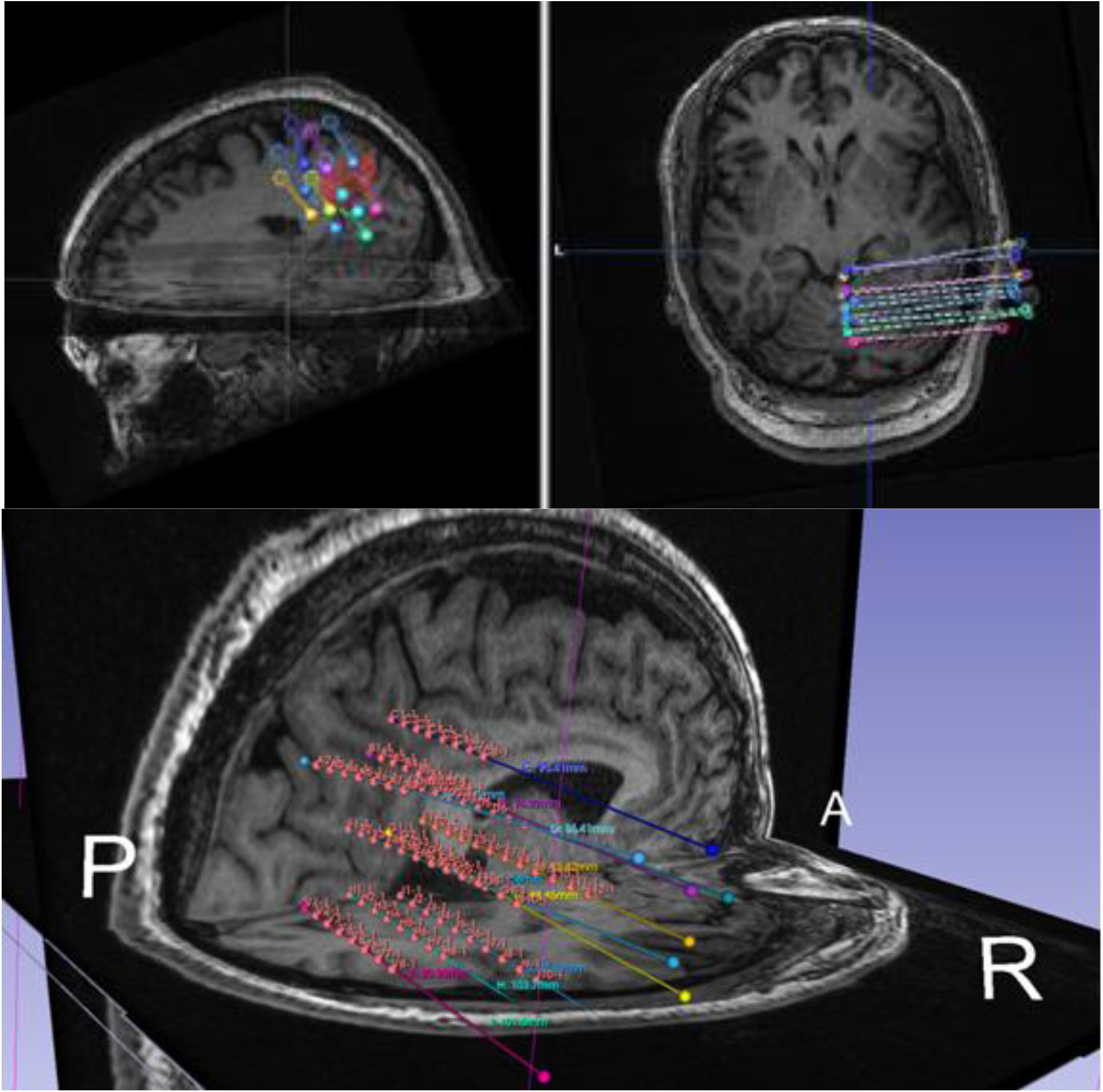
Imaging data from Patient A. Top row: Image received from UMPC showing electrode and lesion locations. **Bottom:** Electrode and lesion location reconstruction in 3D Slicer.

**Supp Figure 3.**
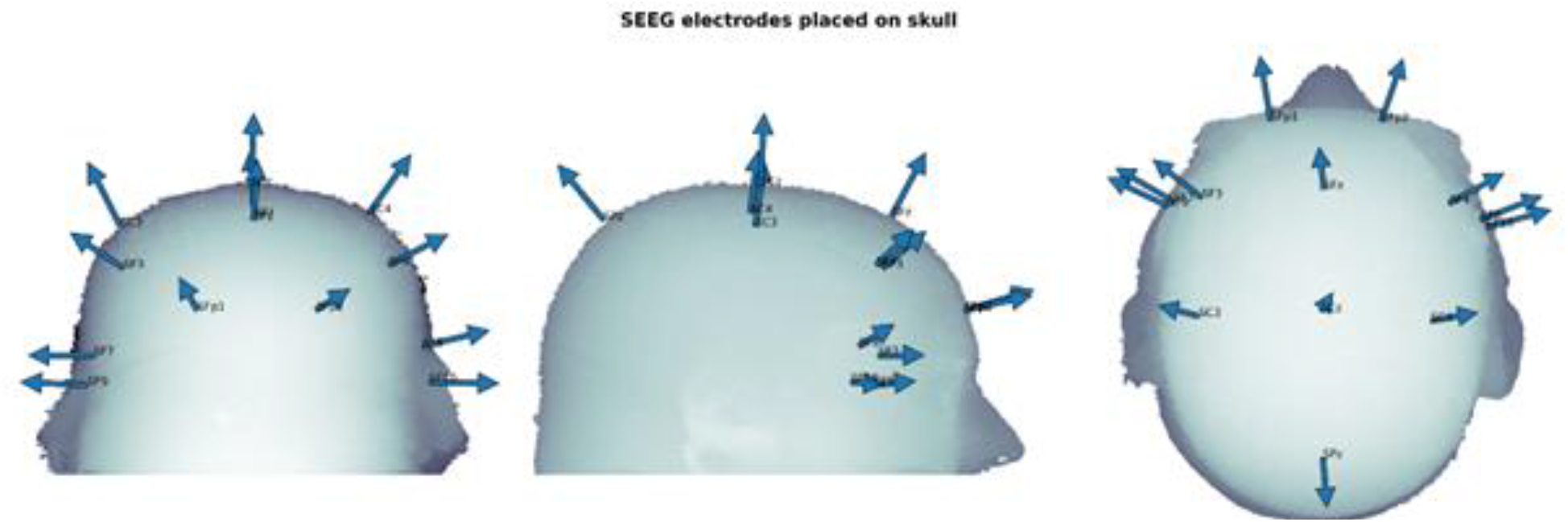
Surface normal vectors originating from the locations of the corresponding EEG electrodes of Patient A placed to a template head model.

**Supp Figure 4.**
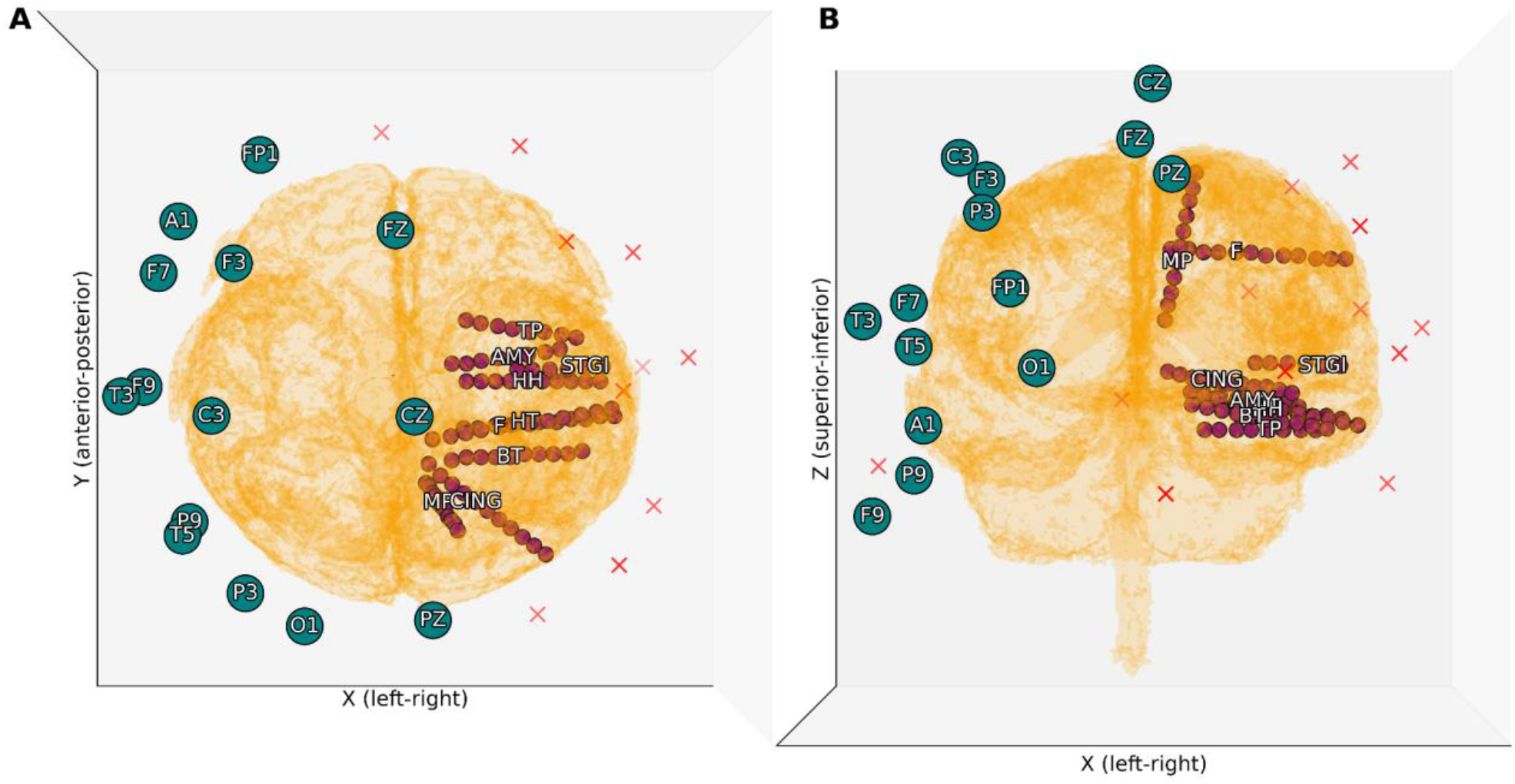
Spatial positioning of the deep and surface electrodes for Patient B. Surface (cap) electrodes with labels are noted in teal dots, missing 10-10 cap electrodes are marked with red crosses, and deep electrodes are marked with purple dots, along with the labels of the shanks. Two different views are shown: superior-inferior (A) and anterior-posterior (B).

## Materials and Methods

**Supp Figure 5.**
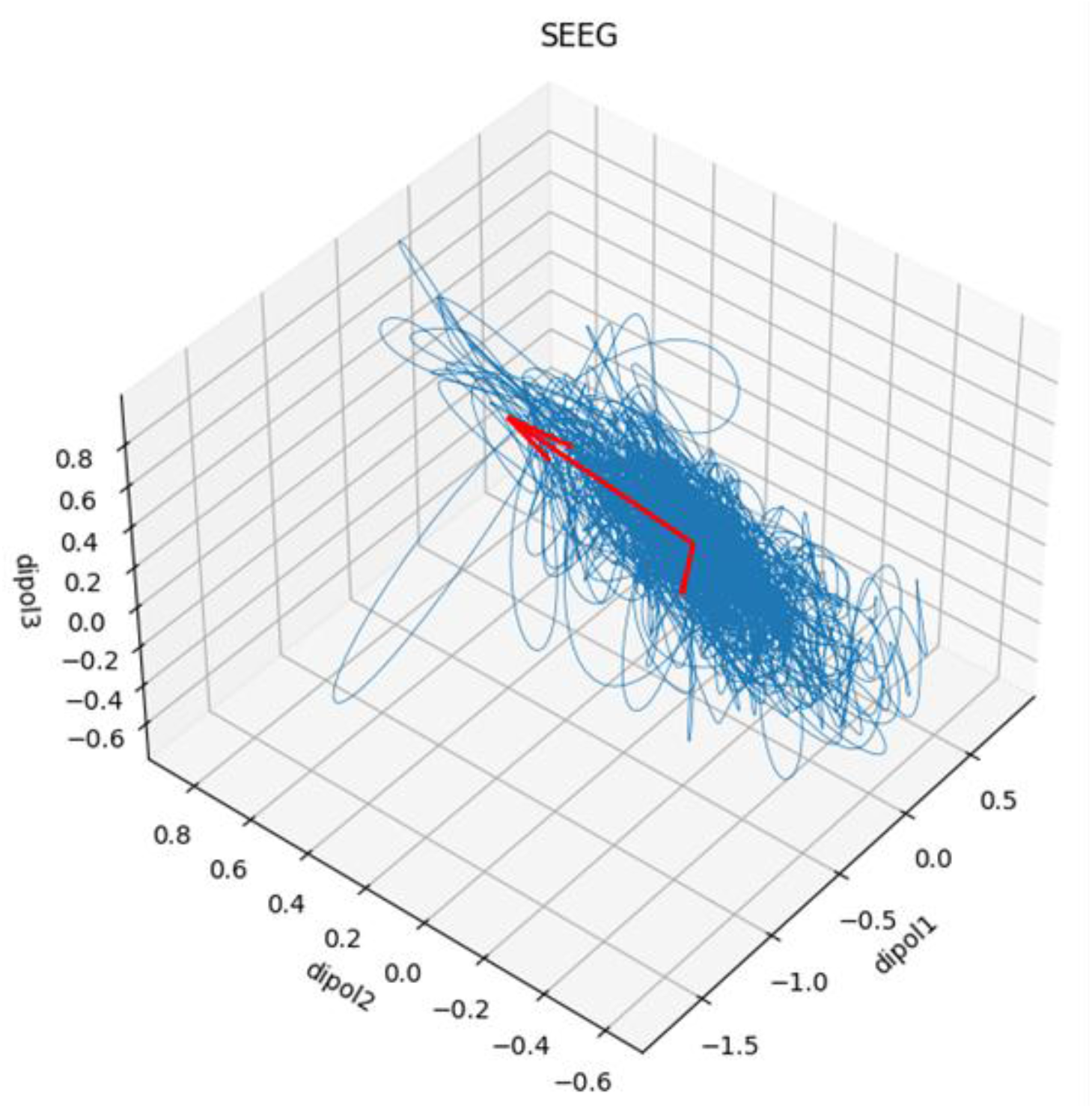
Dipole signal evolution in time. The state of the dipole activity is plotted for every time point in the coordinate space of the dipole components. Red arrows show the orientation and the magnitude of the corresponding eigenvectors (the Principal Component axes) showing a dominant direction.

**Supp Figure 6.**
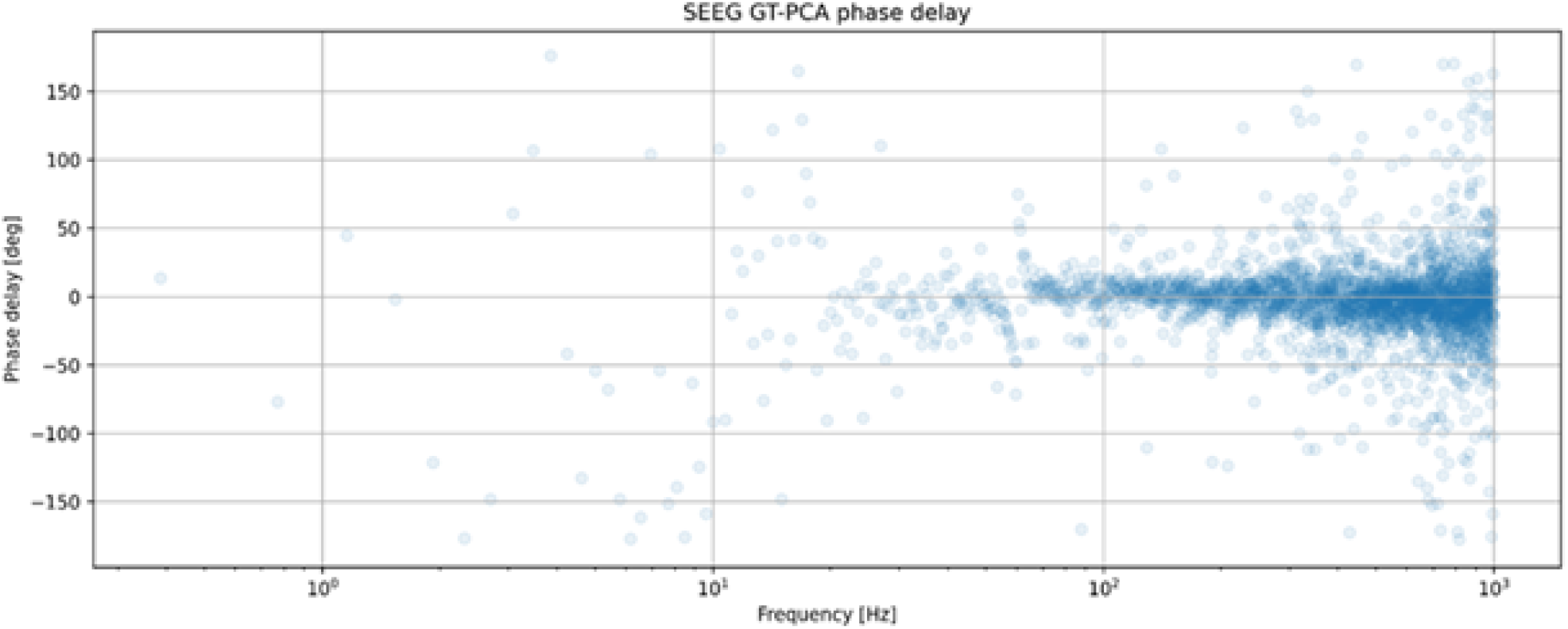
Phase delay function of the recording setup (calculated between the reconstructed data and the SEEG data (ground truth)).

**Supp Figure 7.**
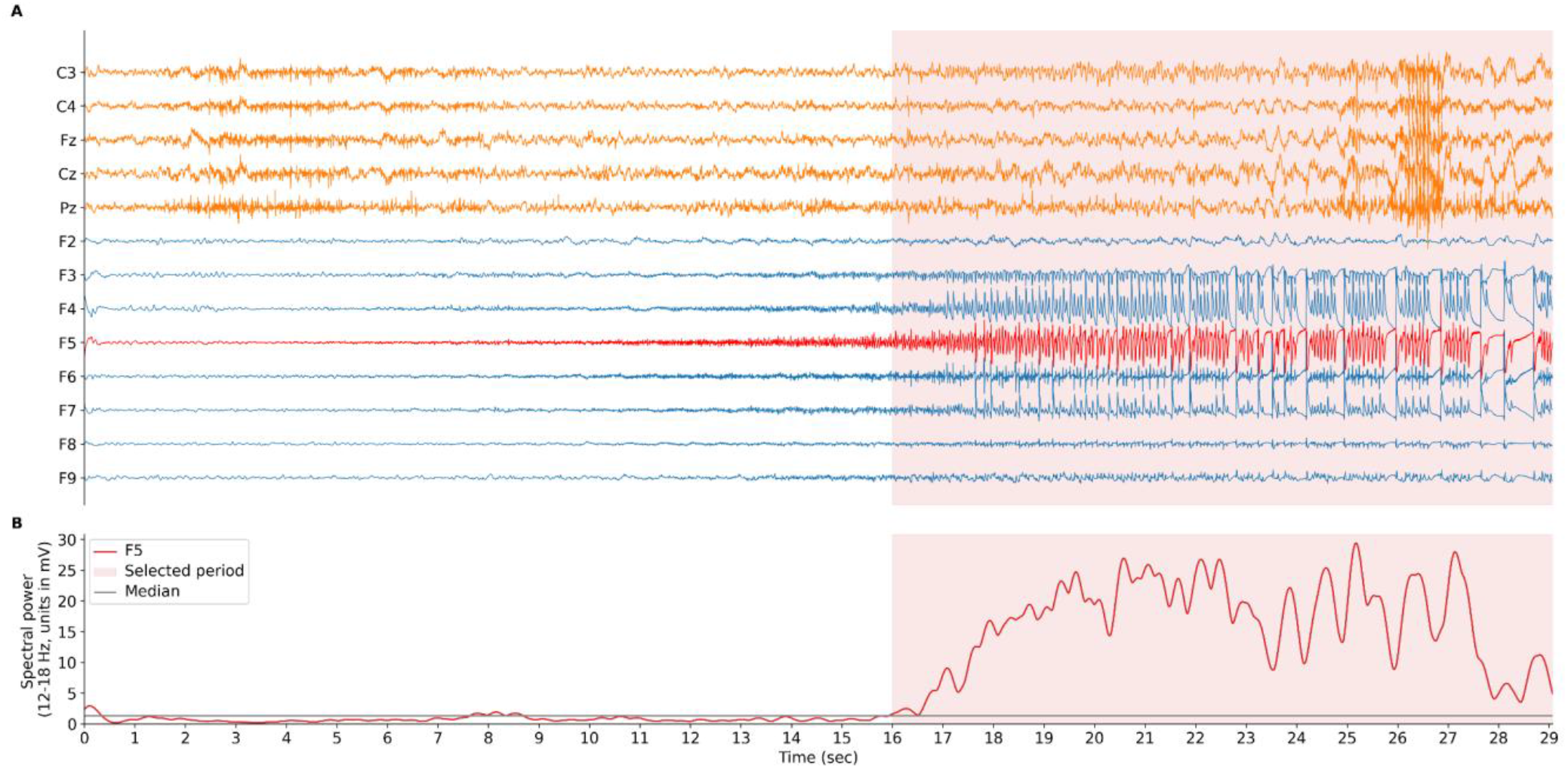
Seizure signals of the analyzed signals for SEEG Patient A. **A:** EEG signals (used for dipole reconstruction) are shown in orange, CSD of intracranial recording shank F is shown in blue, while chosen electrode F5 is highlighted in red. The analyzed period is marked with a red background. **B.** Spectral power of the F5 channel between 12-18 Hz.

Temporal and spectral filtering was based on the prior knowledge of the recording and clinician annotations. The aim was to extract information based on the wavelet transformations of the two channels with the highest power (one from deep and one from surface). Furthermore, deep dipole modeling was necessary to establish a ground truth, despite the lack of information on the position of the SOZ. The 1-45 Hz frequency range during the annotated seizure, and other events is presented on supplementary figure 8A-B. Notations from the EDF file indicate ‘eyes open’ event approximately 10 seconds after the seizure starts and ‘Left facial distortion?’ around 30 seconds. Within these two timestamps, activity in the 2-10 Hz region is visible for both cases, and it is possible that the electromyographic signals did not heavily distort the neural signals. Additionally, the 40-70 second window includes annotations such as ‘screams’, ‘arms straight’ and ‘shaking extremities’. Therefore, despite the high activity visible, we cannot conclude that the similar wavelet patterns observed in this period are solely due to localized neural activity. Furthermore, to highlight our selection of the 10-30 second period of interest, we also calculated the mean Pearson correlation coefficient between all the deep and surface signals in 5 second windows (see supplementary figure 8C). In conclusion, the analyzed signal starts 10 seconds after the original annotation and ends at 30 seconds and is mostly prevalent in the 2-10 Hz frequency range.

